# Peptidergic signaling controls the dynamics of sickness behavior in *Caenorhabditis elegans*

**DOI:** 10.1101/2022.04.16.488560

**Authors:** Javier Marquina-Solis, Elke Vandewyer, Josh Hawk, Daniel A. Colón-Ramos, Isabel Beets, Cornelia I. Bargmann

## Abstract

Pathogenic infection elicits sickness behaviors that promote recovery and survival of the host. For example, following infection with the pathogenic bacterium *Pseudomonas aeruginosa* PA14, the nematode *Caenorhabditis elegans* modifies its sensory preferences to avoid the pathogen. Here we identify antagonistic neuromodulatory circuits that shape this sickness behavior. Using an unbiased cell-directed neuropeptide screen, we show that AVK neurons upregulate and release FMRFamide-like FLP-1 neuropeptides during infection to drive pathogen avoidance. Manipulations that increase or decrease AVK signaling accelerate or delay pathogen avoidance, respectively, implicating AVK in the dynamics of sickness behavior. FLP-1 neuropeptides act via the G-protein-coupled receptor DMSR-7 in RIM/RIC neurons to reduce tyraminergic/octopaminergic signaling that opposes pathogen avoidance. RIM/RIC neurons relay parallel signals from neuropeptides and the cytokine TGF-β that represent internal and external regulators of pathogen avoidance. Our results demonstrate that antagonism between neuromodulatory systems results in slow, graded transitions between alternative behavioral states.

## INTRODUCTION

Animals reorganize their behavioral priorities during pathogenic infection to reduce energy expenditure and facilitate recovery (Aubert et al., 1997; Ayres & Schneider, 2009; Hart, 1988; Hart, 1990; Wang et al., 2016). These behavioral modifications, collectively known as sickness behavior, include lethargy, decreased feeding, and decreased sociability (Dantzer et al., 2008; Maes et al., 2012). Studies of sickness behavior have revealed roles of pro-inflammatory cytokines as conductors of infection signals into the brain (Goehler et al., 1999; Raison et al., 2006; Reyes and Sawchenko, 2002). The neural circuits and mechanisms used by the brain to integrate cytokines and signals from other tissues to coordinate sickness behavior are under active investigation.

Due to its well-defined nervous system and its diverse interactions with bacteria, the nematode *C. elegans* is an excellent model to study sickness behavior. In its natural environment *C. elegans* interacts with many bacterial species, among which are both nutritious and pathogenic *Pseudomonas* species (Dirksen et al., 2016, 2020; Samuel et al., 2016; Schulenburg and Félix, 2017). A well-studied model of *C. elegans* pathogenesis that elicits sickness behavior is the opportunistic pathogenic bacteria *Pseudomonas aeruginosa* PA14 (Laws et al., 2006; Tan et al., 1999a, 1999b; Zhang et al., 2005). *C. elegans* is initially attracted to PA14 and rapidly enters a bacterial lawn to feed. After several hours, PA14 colonizes the *C. elegans* gut and the infected animals exit the PA14 lawn, a sickness behavior termed pathogen avoidance. Failure to avoid PA14 results in death after approximately 72 hours of exposure, while PA14 avoidance prolongs survival (Reddy et al., 2009; Styer et al., 2008). The transition of wild-type worms from an attracted state to an avoiding state represents a well-defined natural sickness behavior.

Previous studies have identified several components of sickness behavior elicited by sensory cues associated with PA14 (Meisel et al., 2014; Reddy et al., 2009; Styer et al., 2008). The ASJ chemosensory neurons contribute to PA14 avoidance by recognizing the secondary bacterial metabolites phenazine-1-carboxamide (PCN) and pyochelin, and secreting the cytokine Transforming Growth Factor beta (TGF-β)/DAF-7 (Meisel et al., 2014). TGF-β signaling activates PA14 avoidance by suppressing animals’ preference for the low oxygen levels in a PA14 lawn (Meisel et al., 2014). Conversely, *npr-1* mutants, which have enhanced low oxygen preference, fail to avoid PA14 (Reddy et al., 2009; Styer et al., 2008). Reducing environmental oxygen restores PA14 avoidance to infected *daf-7* and *npr-1* mutants, suggesting that oxygen preferences interact with a second system that recognizes infection to alter behavior (Meisel et al., 2014).

While the sensory cues PCN, pyochelin, and oxygen are present throughout PA14 exposure, PA14 avoidance increases after several hours of pathogenic infection through internal detection of pathogenesis. Here we describe a peptidergic signaling pathway that sets the tempo of the slow avoidance of pathogenic bacteria. Using genetic and molecular tools, we demonstrate that AVK interneurons drive PA14 avoidance by release of FMRFamide-like neuropeptides encoded by *flp-1*. FLP-1 neuropeptides are detected by tyraminergic and octopaminergic neurons that oppose pathogen avoidance. Transcriptional regulation within multiple neurons integrates sensory and physiological information that supports sickness behavior.

## RESULTS

### Neuropeptide processing affects pathogen avoidance

To gain insight into molecular mechanisms that drive pathogen avoidance, we examined 67 mutants affecting neurotransmitters, neuropeptides, innate immunity and stress response in a PA14 avoidance assay **(Table S1)**. For the initial screen, we placed L4 stage animals on a small lawn of PA14 bacteria and calculated the fraction of adult animals outside the lawn after 20 hours (Avoidance Ratio) (**Figure 1A**). Over 90% of wild-type animals were outside the PA14 lawn after 20 hours (**Figure 1A**), but this behavior was greatly reduced in *egl-3* mutants. *egl-3* encodes a Proprotein Convertase type 2 (PC2), an enzyme that converts pro-peptides into neuropeptide intermediates (Husson et al., 2006) (**Figure 1B**). Two *egl-3* loss-of-function mutants showed strong PA14 avoidance defects such that less than 15% of animals avoided PA14 at 20 hours (**Figure 1C**). A second gene affecting neuropeptide processing, *egl-21*, encodes a Carboxypeptidase E (CPE) that releases C-terminal basic amino acids from neuropeptide precursors (Husson et al., 2007). Like *egl-3* mutants, *egl-21* loss-of-function mutants showed minimal PA14 avoidance (**Figure 1C**). These results suggest that PA14 avoidance is modulated by neuropeptides.

**Figure 1.**
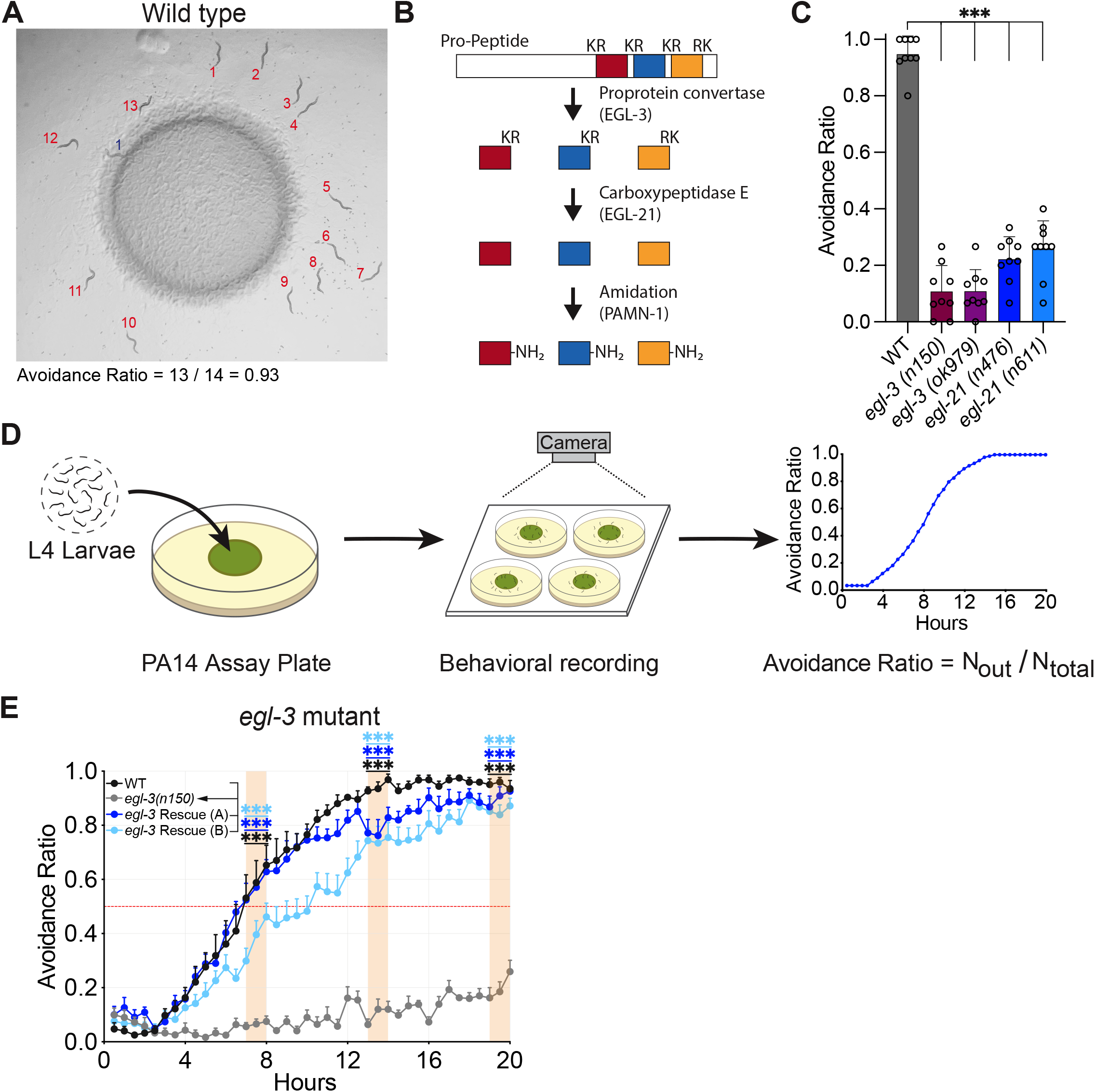
Neuropeptide processing enzymes are required for PA14 avoidance. **A**, Wild-type animals after 20 hours exposure to a *Pseudomonas aeruginosa* PA14 lawn. Red numbers indicate animals outside the lawn, blue number indicates animal inside the lawn. **B**, Schematic of neuropeptide processing. **C**, Avoidance Ratio of wild-type animals and *egl-3(n150)*, *egl-3(ok979)*, *egl-21(n476)*, and *egl-21(n611)* loss-of-function mutants calculated 20 hours after exposure to PA14. **D**, Schematic of dynamic PA14 avoidance assay. In all figures, a single assay represents the behavior of 11-17 animals on a single plate. **E**, The PA14-avoidance defect of *egl-3(n150)* mutants can be rescued by expression of a genomic fragment containing the *egl-3* gene. All figures show comparisons of average Avoidance Ratio at hours 7 to 8, 13 to 14, and 19 to 20 after PA14 exposure, using test animals and controls tested in parallel on the same day. For detailed statistics see **Table S4**. For experiments in **C,** *n* = 9 assays for all groups; for experiments in **E**, *n* = 8 assays for all groups. Graphs are mean + s.e.m. ***P < 0.001, by one-way repeated-measures analysis of variance (ANOVA) with Dunnett’s post-hoc test; (**C**) comparisons to WT, (**E**) comparisons to *egl-3(n150)*.

To gain insight into the dynamics of PA14 avoidance, individual assays were video recorded and the Avoidance Ratio was calculated at 30 minute intervals (**Figure 1D**). We quantified and compared the average Avoidance Ratio for each group at hours 7 to 8, 13 to 14, and 19 to 20. Wild-type animals started to avoid PA14 by 4 hours after exposure, gradually increasing their avoidance until 95% of animals were outside the PA14 lawn (**Figure 1E**); *egl-3* mutants were defective in PA14 avoidance at all times. Two transgenic strains expressing an *egl-3* genomic fragment in the *egl-3(n150)* mutant background exhibited normal PA14 avoidance behavior, confirming that the mutant phenotype was caused by loss of *egl-3* function (**Figure 1E**).

### The peptidergic AVK neurons mediate PA14 pathogen avoidance

Neuropeptides are produced by most of the 302 neurons in the *C. elegans* nervous system as well as the intestine and other non-neuronal cells (Taylor et al., 2021). To identify cells that modulate PA14 avoidance, we used CRISPR/Cas9 to insert LoxP sites flanking the endogenous *egl-3* locus and expressed Cre recombinase in selected cells to inactivate *egl-3* (**Figure 2A**). Animals carrying the *egl-3* floxed allele resembled wild-type animals in the PA14 avoidance assay **(Figure S1B)**. Pan-neuronal *egl-3* knockout diminished PA14 avoidance, while intestinal *egl-3* knockout did not affect the behavior **(Figures S1C and S1D)**. Next, we examined 25 transgenic lines expressing Cre under different neuronal promoters and quantified PA14 avoidance at 20 hours **(Figure S1A and Table S2)**. Remarkably, knockout of *egl-3* only in the peptidergic AVK neurons, using two different cell-specific promoters, was sufficient to recapitulate the *egl-3* mutant defect (**Figures 2B and S1E)**. This result suggests that the *egl-3* defect can largely be explained by loss of neuropeptides from the AVK neurons, although we did not test all neurons in this screen.

**Figure 2.**
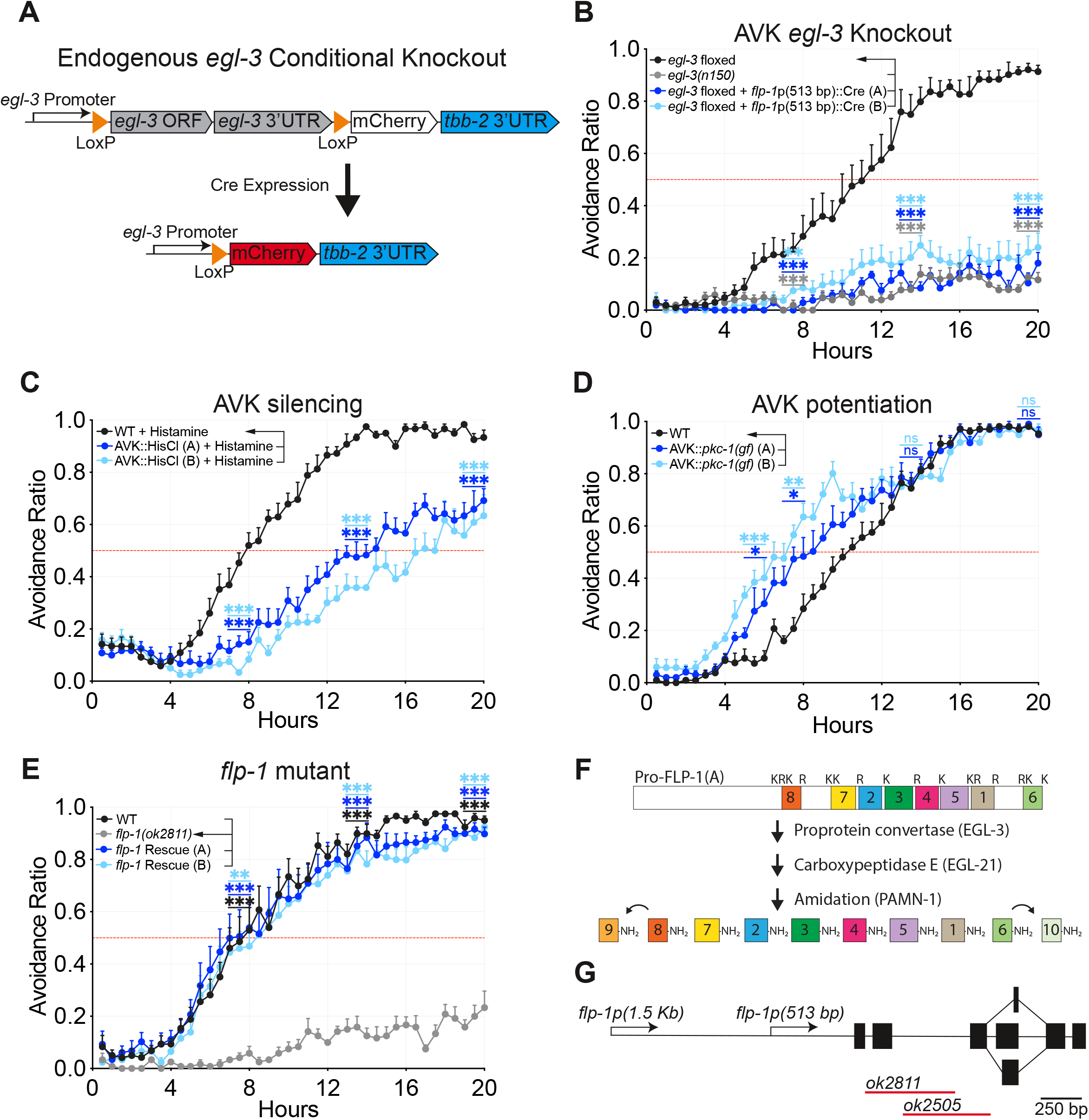
The AVK neurons promote PA14 avoidance via FLP-1 neuropeptides. **A**, Schematic of endogenous *egl-3* knockout strategy using Cre-lox recombination. In the *egl-3-*floxed strain, expression of Cre recombinase under cell-specific promoters excises the *egl-3* gene, allowing mCherry expression in the targeted cells. **B**, Cell-specific knockout of *egl-3* using the *flp-1*(513 bp) promoter to drive Cre only in the AVK neurons; see also **Figure S1E**. **C**, Silencing of the AVK neurons by cell-specific expression of the histamine-gated chloride channel HisCl1; PA14 plates were supplemented with 10 mM histamine. **D**, Potentiation of dense core vesicle release from the AVK neurons by cell-specific expression of a constitutively active protein kinase C (*pkc-1(gf)*). **E**, *flp-1(ok2811)* mutants are defective in PA14 avoidance and rescued by transgenic expression of a *flp-1* genomic fragment. **F**, Schematic of the Pro-FLP-1-A precursor and the proteolytic pathway that yields active FLP-1 peptides. FLP-1-8 and FLP-1-9 are RYamide peptides. FLP-1-9 and FLP-1-10 are derived from FLP-1-8 and FLP-1-6 respectively (Van Bael et al., 2018). **G**, Schematic of the *flp-1* gene structure depicting 5’-regulatory regions, exons and splice isoforms. The *ok2811* null allele contains a deletion that disrupts the *flp-1* reading frame. The deletion in *ok2505* is a partial loss-of-function that results in the loss of FLP-1 RYamide peptides KPNFMRYa and PNFMRYa, and the FLP-1 RFamide peptide AGSDPNFLRFa (Buntschuh et al., 2018). For experiments in **B**, **C**, and **E**, *n* = 8 assays for all groups; for experiments in **D**, *n* = 7 assays for all groups. Graphs are mean + s.e.m. *P < 0.05, **P < 0.01, ***P < 0.001, ns, not significant, by one-way ANOVA with Dunnett’s post-hoc test; (**B**, **E**) comparisons to *egl-3* floxed, (**C**, **D**) comparisons to WT.

To ask if AVK acts acutely during PA14 avoidance, we expressed the *Drosophila* histamine-gated chloride channel HisCl1 in the AVK neurons. *C. elegans* does not use histamine as a neurotransmitter, but takes it up from the environment, allowing silencing of AVK::HisCl1 neurons by acute histamine addition (Pokala et al., 2014). The time at which 50% of animals avoided PA14 was delayed by 5 to 8 hours in animals in which AVK was silenced, and the avoidance plateau was reduced by ∼25% (**Figure 2C**). In control experiments performed in the absence of histamine, PA14 avoidance of wild-type animals and AVK::HisCl1 animals was indistinguishable **(Figure S2A)**. In a second experiment, we expressed the tetanus toxin light chain, which cleaves synaptobrevin protein to inhibit synaptic and peptidergic vesicle release, in AVK. AVK::Tetanus Toxin animals had a similar delay in PA14 avoidance as AVK::HisCl1 animals **(Figure S2B)**. These experiments demonstrate that AVK activity and AVK vesicle release promote PA14 avoidance.

As a complementary experiment, we expressed a constitutively active Protein Kinase C (*pkc-1*(*gf*)) in the AVK neurons. *pkc-1(gf)* potentiates release of neuropeptidergic dense core vesicles and synaptic vesicles in *C. elegans* (Sieburth et al., 2007); in the peptidergic AVK neurons, it is predicted to potentiate neuropeptide release. The time at which 50% of animals avoided PA14 was accelerated by ∼4 hours in the AVK::*pkc-1*(*gf*) strain (**Figure 2D**). AVK::*pkc-1*(*gf*) animals did not avoid OP50, a non-pathogenic *E. coli* strain use as the standard laboratory food source **(Figure S2C)**. Together, the inactivation experiments and gain of function experiments indicate that AVK neuropeptide secretion is rate-limiting for PA14 avoidance.

### FMRFamide-like peptides are released from AVK neurons during infection

To further characterize the relationship between neuropeptide processing and AVK, we examined two neuropeptides that are highly enriched and expressed at high levels in AVK: *flp-1* and *nlp-49* (Lorenzo et al., 2020; Taylor et al., 2021)*. flp-1(ok2811)* loss-of-function mutants had a defect that closely resembled that of *egl-3* AVK-specific knockouts (**Figure 2E**), whereas *nlp-49* mutants were normal in PA14 avoidance **(Figure S3A)**. The PA14 avoidance defect in *flp-1(ok2811)* was rescued by a *flp-1* genomic fragment that was expressed exclusively in AVK (**Figure 2E, Figure S3C).** Although *flp-1* can be expressed in AIY as well as AVK neurons (Jia and Sieburth, 2021; Nelson et al., 1998), the reporters and transgenes studied here appeared to be AVK-specific **(Methods)**.

The *flp-1* gene encodes eight neuropeptides with a C-terminal RFamide sequence and two neuropeptides with a C-terminal RYamide sequence (Van Bael et al., 2018; Husson et al., 2005; Rosoff et al., 1993) (**Figure 2F and Table S3)**. *flp-1(ok2505)* mutants, which lack FLP-1 RYamide-peptides (Buntschuh et al., 2018), showed normal PA14 avoidance, suggesting that the FLP-1 RFamide-peptides are sufficient for the behavior (**Figures 2G and S3B)**.

To characterize the relationship between *flp-1* and pathogenesis, we compared the susceptibility of wild type and *flp-1* mutants to killing by PA14. On standard PA14 assay plates, *flp-1* mutants were more susceptible to PA14 killing than wild-type animals, with a t_50_ of killing at ∼60 hours instead of ∼96 hours (**Figure 3A**). To ask whether this susceptibility arises from reduced PA14 avoidance, killing rates were examined on plates uniformly seeded with PA14, so neither *flp-1* mutants nor wild-type animals could avoid the pathogen. Under these conditions, both wild type and *flp-1* mutants had a t_50_ of killing at ∼60 hours (**Figure 3B**). These results indicate that the increased susceptibility of *flp-1* mutants to killing by PA14 can be explained by their failure to avoid the bacterial pathogen.

**Figure 3.**
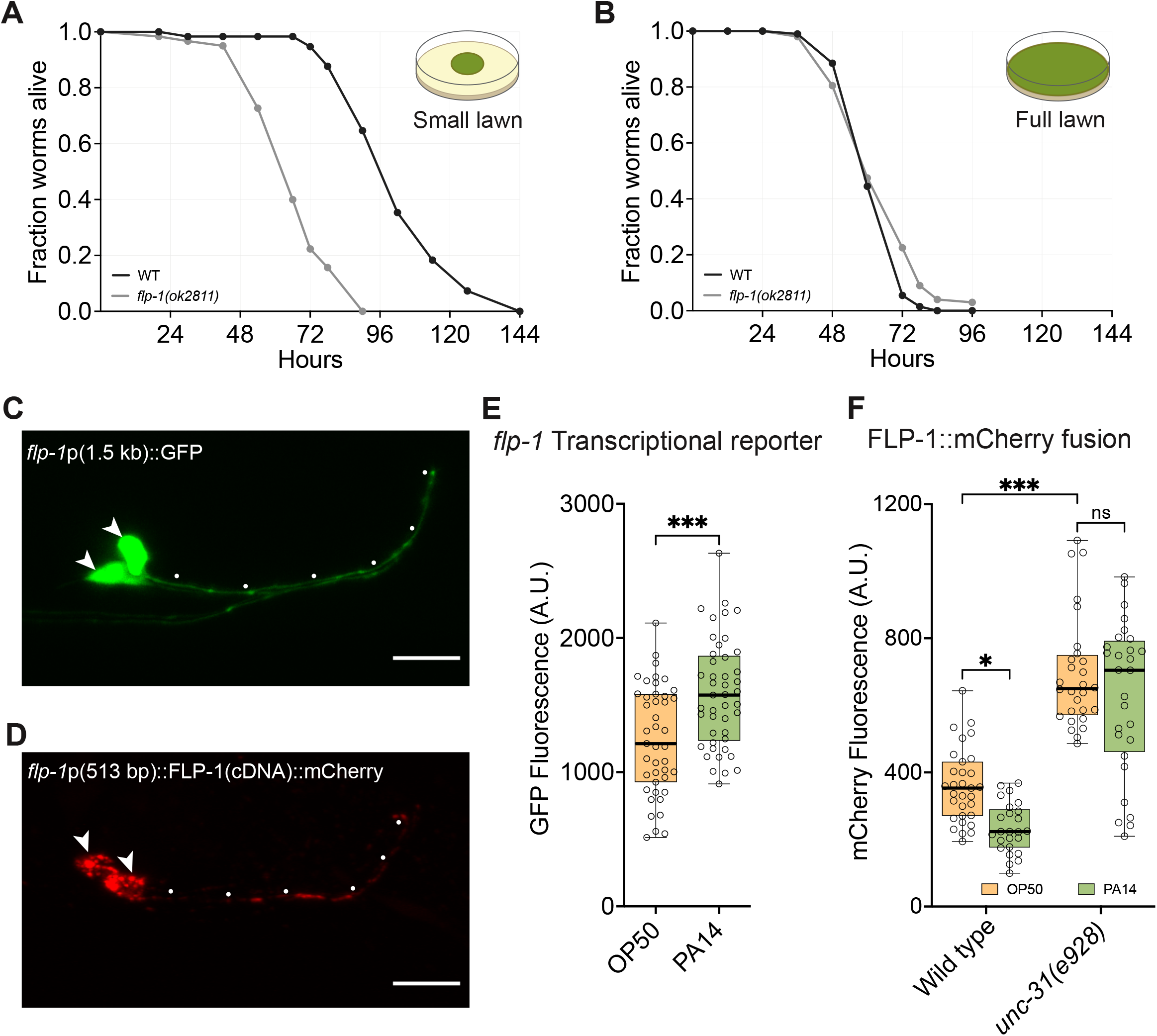
FLP-1 neuropeptides are released during exposure to PA14. **A**, **B**, Fraction alive over time for WT and *flp-1(ok2811)* mutants on small PA14 lawns (**A**), and full PA14 lawns (**B**). Graphs are means from 3 (**A**) and 2 (**B**) independent experiments. **C**, **D**, Images showing AVK neurons expressing GFP under the *flp-1*(1.5 kb) promoter (**C**) and expressing FLP-1(cDNA)::mCherry under the *flp-1*(513 bp) promoter (**D**). Arrowheads indicate AVK cell bodies; dots indicate AVK axons. Anterior at right, dorsal up. Scale bars, 10 μm. **E**, Animals expressing *flp-1*p(1.5 kb)::GFP exposed to *E. coli* OP50 or full PA14 lawns; exposure to PA14 upregulates the transcriptional reporter. **F**, Animals expressing *flp-1*p(513 bp)::FLP-1(cDNA)::mCherry exposed to OP50 or full PA14 lawns; in wild-type animals (left) exposure to PA14 decreases FLP-1(cDNA)::mCherry levels in the AVK cell body. *unc-31(e928)* mutants in OP50 have higher FLP-1(cDNA)::mCherry levels in the AVK cell bodies compared to wild-type, with no significant difference between *unc-31(e928)* mutants in OP50 or PA14, suggesting that *unc-31*-dependent dense core vesicle release drives the decrease in FLP-1(cDNA)::mCherry in PA14 animals. For experiments in **E** and **F**, box plots’ center line denotes median; box range denotes 25–75th percentiles; whiskers denote minimum–maximum values. For experiments in **E**, *flp-1*p(1.5 kb)::GFP in OP50, *n* = 43; in PA14, *n* = 45. For experiments in **F**, WT *flp-1*p(513 bp)::FLP-1(cDNA)::mCherry in OP50, *n* = 31, in PA14, *n* = 28; *unc-31(e928) flp-1*p(513 bp)::FLP-1(cDNA)::mCherry in OP50, *n* = 25, in PA14, *n* = 28. *P < 0.05, ***P < 0.001, ns, not significant, (**E**) by unpaired two-tailed *t*-test, or (**F**) by two-way ANOVA with Tukey’s post-hoc test.

To understand how pathogen infection affects *flp-1* activity, we examined a transcriptional reporter in which a 1.5 kb fragment upstream of *flp-1* drove GFP expression. This *flp-1* reporter gene was upregulated in AVK within 4 hours of exposure to PA14, suggesting that PA14 infection increases FLP-1 biosynthesis (**Figures 3C and 3E**).

We next used a FLP-1::mCherry fusion protein to monitor FLP-1 levels in AVK neurons as a proxy for neuropeptide release (Oranth et al., 2018) (**Figure 3D**). In this experiment, FLP-1::mCherry was expressed from a 513 bp *flp-1* promoter element that is specific to AVK but not significantly induced by PA14 infection **(Figure S3D)**. Animals exposed to PA14 for 4 hours had a ∼50% decrease in FLP-1::mCherry fluorescence compared to animals in *E. coli* OP50, consistent with increased release of FLP-1 neuropeptides in PA14 (**Figure 3F**). To establish that this decrease in FLP-1::mCherry results from dense core vesicle exocytosis, we examined mutants lacking *unc-31*/CAPS (Ca^2+^-activated protein for secretion), which are deficient in neuropeptide secretion (Ann et al., 1997; Speese et al., 2007). *unc-31* mutants had increased FLP-1::mCherry fluorescence, consistent with reduced neuropeptide release, and showed no difference in FLP-1::mCherry fluorescence after exposure to PA14 (**Figure 3F).** These results suggest that FLP-1::mCherry release through dense core vesicle exocytosis is enhanced by PA14 infection.

### FLP-1 peptides signal through the G protein-coupled receptor DMSR-7

The *C. elegans* genome contains 158 neuropeptide precursor genes, encoding over 250 distinct neuropeptides, and more than 100 neuropeptide G-protein-coupled receptor (GPCR) genes (Frooninckx et al., 2012; Li and Kim, 2010). Previous screens of neuropeptide receptors using a heterologous expression system identified FRPR-7, NPR-4, NPR-5, NPR-6, and NPR-11 as receptors for FLP-1 neuropeptides (Kubiak et al., 2008; Lowery et al., 2003; Oranth et al., 2018). FLP-1 neuropeptides released from AVK regulates acute locomotion parameters such as speed and body posture through the activation of NPR-6 and FRPR-7 receptors on motor neurons (Oranth et al., 2018). Although *npr-6(tm1497) frpr-7(gk463846)* double mutants had defects in body posture and locomotion, their PA14 avoidance was normal **(Figure S4A)**. Loss-of-function mutants of *npr-4*, *npr-5*, and *npr-11* did not affect PA14 avoidance **(Table S1)**.

To identify additional FLP-1 receptors, we probed a library of *C. elegans* G-protein coupled neuropeptide receptors in CHO cells expressing the promiscuous G-protein subunit G_α16_ and mitochondrially targeted aequorin (Janssen et al., 2008). Activation of the receptor in this CHO cell line evokes Ca^2+^ increase and luminescence. Cells transfected with a *dmsr-7* cDNA construct generated Ca^2+^ responses upon exposure of FLP-1 RFamide peptides at concentrations of 0.1–3 nM (**Figure 4A and Table S3)** and responded to other families of FMRFamide-like peptides at higher concentrations **(Vandewyer and Beets, in preparation)**. DMSR-7 is the highest-potency FLP-1 receptor we have identified. We also identified DMSR-1, DMSR-5 and DMSR-6 as receptors that respond to FLP-1 RFamide peptides **(Vandewyer and Beets, in preparation)**.

**Figure 4.**
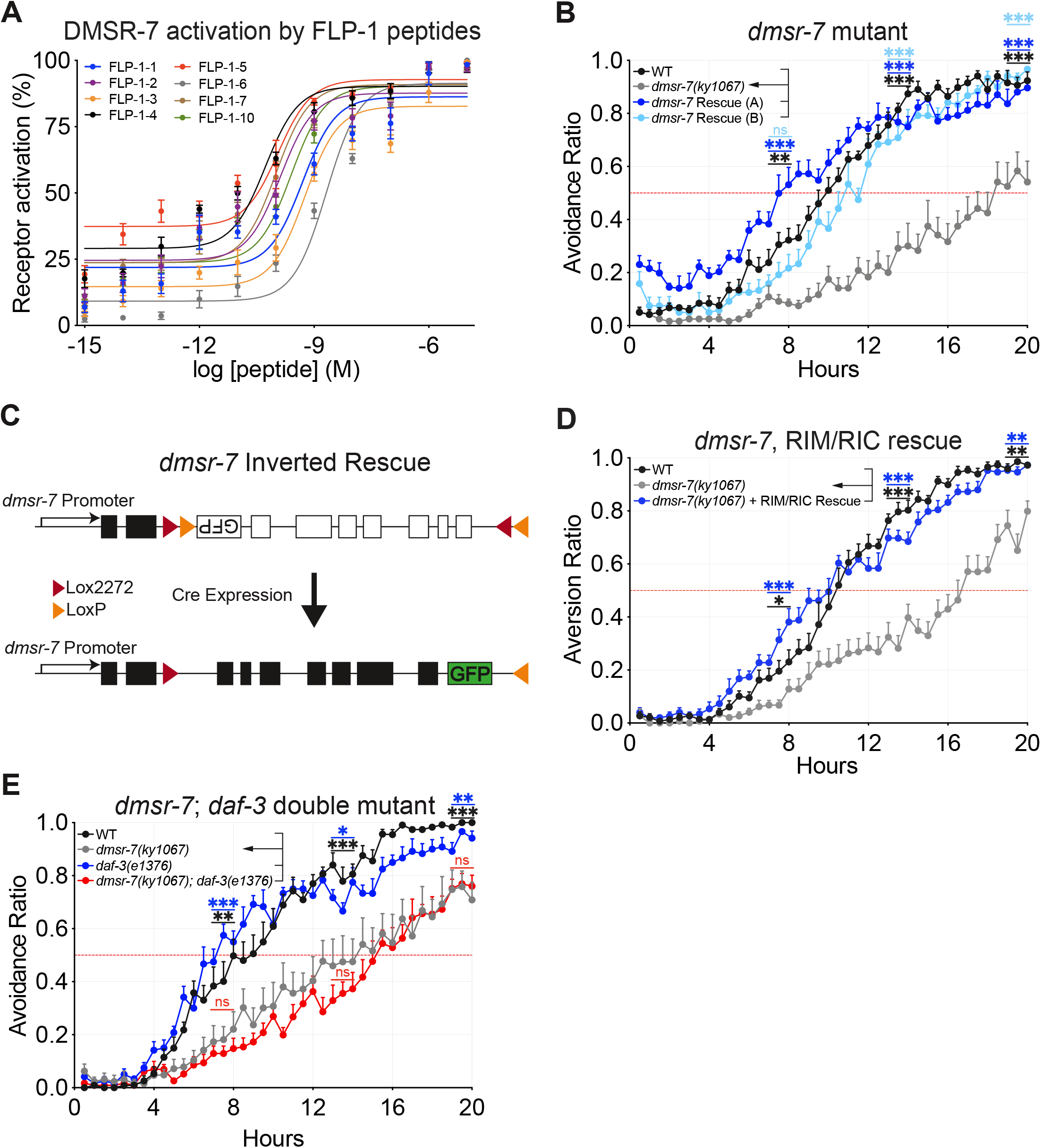
FLP-1 neuropeptides signal through the DMSR-7 receptor. **A**, Aequorin bioluminescence (% of maximum receptor activation) of CHO cells expressing Gα16 and DMSR-7 in response to different concentrations of FLP-1 peptides (#1-7, 10). **B**, Rescue of the *dmsr-7(ky1067)* mutants by transgenic expression of *dmsr-7* from its endogenous promoter. **C**, Schematic of the transgene used to rescue *dmsr-7* in subsets of its endogenous expression pattern using the Cre-lox system. Reconstitution of the transgene is confirmed by GFP expressed with *dmsr-7* in a bicistronic transcript. **D**, Rescue of *dmsr-7* in the RIM and RIC interneurons restores PA14 avoidance to wild-type levels. Expression of *dmsr-7* in a set of interneurons and motor neurons not including RIM or RIC also resulted in significant rescue of *dmsr-7(ky1067)*, suggesting that *dmsr-7* may act in additional cell types to regulate pathogen avoidance (Figure S5E). **E**, *dmsr-7 daf-3* double mutants behave like *dmsr-7* single mutants in the PA14 avoidance assay. For experiments in **A**, *n* = 6 assays for all groups; for experiments in **B**, **E**, *n* = 8 assays for all groups; for experiments in **D**, *n* = 10 assays for all groups. Graphs are mean + s.e.m. *P < 0.05, **P < 0.01, ***P < 0.001, ns, not significant, (**B**, **D**, **E**) by one-way ANOVA with Dunnett’s post-hoc test; comparisons to *dmsr-7(ky1067)*.

Examining loss-of-function alleles for each FLP-1 receptor candidate revealed that *dmsr-7(ky1067)* mutants were defective in PA14 avoidance, while *dmsr-1(ky1053)*, *dmsr-5(ky1059)* and *dmsr-6(ky1065)* mutants were normal (**Figures 4B and S4B-D)**. PA14 avoidance in *dmsr-7(ky1067)* mutants was rescued by a transgene encompassing the *dmsr-7* genomic region, confirming that the defect was caused by loss of *dmsr-7* (**Figure 4B**). *dmsr-7* mutants were less defective in PA14 avoidance than *flp-1* mutants **(Figure S4F)**, suggesting the involvement of additional receptors. We tested *dmsr-1(ky1053) dmsr-7(ky1067)* double mutants and *dmsr-5(ky1059) dmsr-6(ky1065) dmsr-7(ky1067)* triple mutants for PA14 avoidance but found no difference from *dmsr-7* single mutants **(Figures S4E and S4F)**.

### FLP-1 and TGF-β signals converge on RIM and RIC interneurons

The *dmsr-7* genomic fragment that rescued PA14 avoidance in *dmsr-7* mutants is expressed in motor neurons and interneurons but not in sensory neurons **(Figure S5A)**. To identify neurons in which DMSR-7 acts to induce PA14 avoidance, we used an inverted Cre-LoxP intersectional strategy to rescue *dmsr-7(ky1067)* in subsets of its endogenous expression pattern (**Figure 4C**). Control experiments showed that the inverted *dmsr-7* transgene did not rescue *dmsr-7(ky1067)* mutants, but PA14 avoidance was fully rescued upon pan-neuronal Cre expression **(Figures S5B and S5C)**. By limiting Cre expression to neuronal subsets, we observed full rescue when *dmsr-7* was expressed only in the RIM and RIC interneurons and significant rescue when *dmsr-7* was expressed only in RIC (**Figures 4D and S5D)**.

Previous work demonstrated that the RIM and RIC interneurons contribute to PA14 avoidance behavior (Meisel et al., 2014). When ASJ chemosensory neurons detect the PA14 secondary-metabolites PCN and pyochelin, they induce transcription of the cytokine DAF-7/TGF-β. DAF-7 activates the DAF-1/DAF-4 TGF-β receptor in RIM and RIC neurons, resulting in the inhibition of the co-SMAD transcription factor, DAF-3. Loss-of-function mutants of *daf-7* or *daf-1* display severe defects in PA14 avoidance that are fully suppressed by *daf-3* (Meisel et al., 2014) **(Figures S6A and S6B)**.

Because *daf-7(e1372)* and *flp-1(ok2811)* mutants displayed similar phenotypes on the PA14 avoidance assay and the two signaling pathways converged at the RIM and RIC interneurons, we sought to understand their relationship. *daf-3(e1376)* fully suppressed the *daf-7(e1372)* PA14 avoidance defect, but it did not suppress *flp-1(ok2811)* or *dmsr-7(ky1067)* defects (**Figures 4E, S6A, and S6C)**. Thus, while FLP-1 and DAF-7 signals converge in RIM and RIC neurons, they promote PA14 avoidance by distinct mechanisms.

### *dmsr-7* opposes tyraminergic/octopaminergic signaling from RIM/RIC neurons

DMSR-7 is related to the *Drosophila* Myosuppressin receptor 1, which signals through an adenylate cyclase-inhibiting GPCR pathway (Egerod et al., 2003). By analogy, DMSR-7 might inhibit RIM and RIC interneurons during exposure to PA14 (**Figure 5A**). In agreement with this hypothesis, expressing the tetanus toxin light chain to block neurotransmitter release from RIM and RIC resulted in full restoration of PA14 avoidance to *dmsr-7(ky1067)* mutants (**Figure 5B**). Expressing tetanus toxin in RIM and RIC in wild-type animals had minimal effects on PA14 avoidance. These results suggest that in *dmsr-7(ky1067)* mutants, synaptic release from RIM and RIC delays PA14 avoidance, and FLP-1 signaling through DMSR-7 opposes this function.

**Figure 5.**
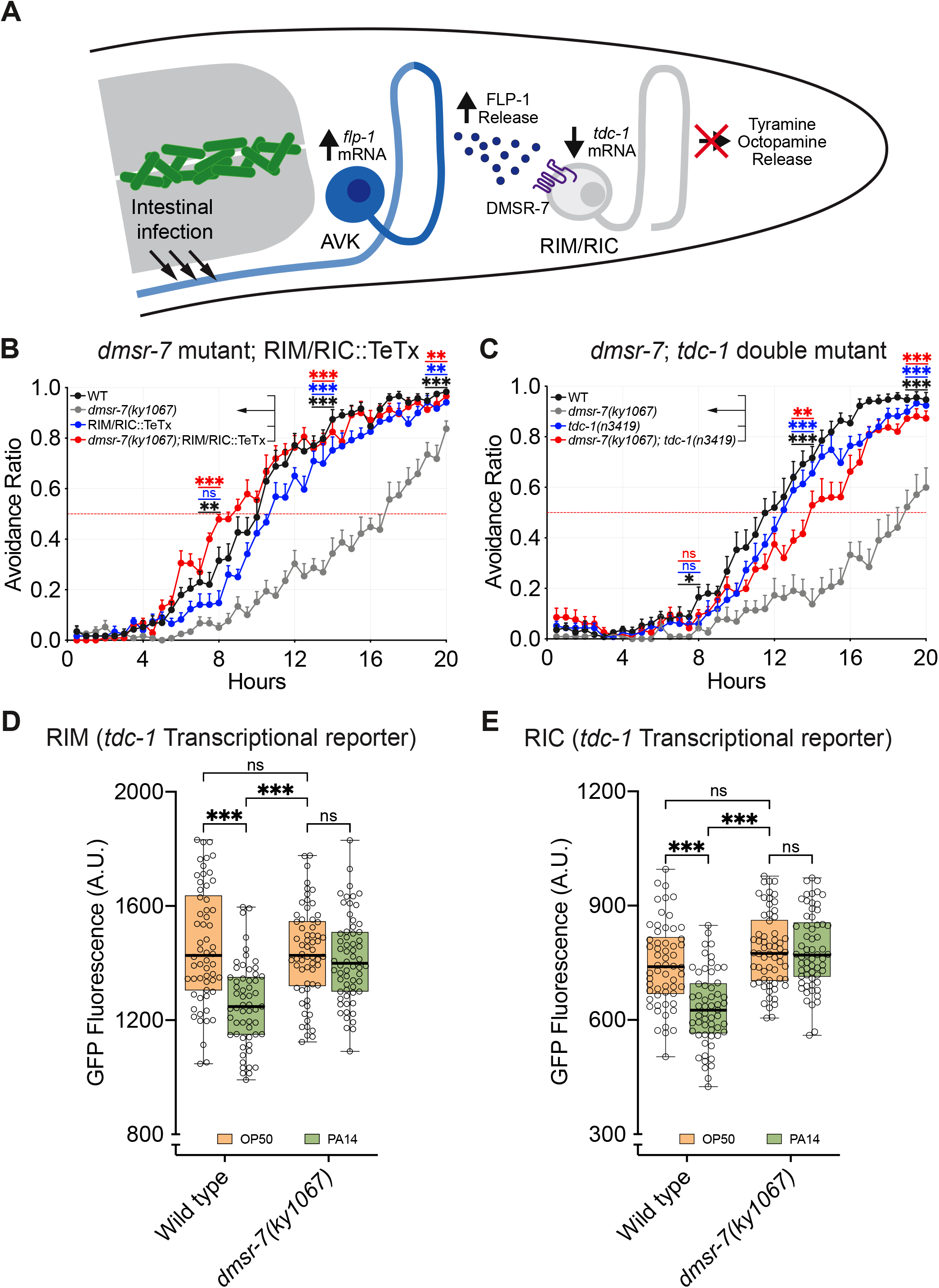
FLP-1 signaling opposes tyraminergic and octopaminergic signaling. **A**, Model: the AVK neurons release FLP-1 neuropeptides during exposure to PA14, suppressing tyraminergic and octopaminergic signaling to promote pathogen avoidance. **B**, Expression of tetanus toxin light chain in RIM and RIC interneurons blocks neurotransmitter release and suppresses the *dmsr-7* PA14-avoidance defect. **C**, *tdc-1* encodes a tyrosine decarboxylase required for the synthesis of both tyramine and octopamine. *tdc-1(n3419)* suppresses the avoidance defect of *dmsr-7(ky1067)* mutants. **D**, **E**, PA14 exposure downregulates a *tdc-1*p::GFP reporter in a *dmsr-7*-dependent manner in RIM(**D**) and RIC(**E**). For experiments in **B** and **C**, *n* = 8 assays for all groups; for experiments in **D**, **E**, WT in OP50, *n* = 58; WT in PA14, *n* = 57; *dmsr-7* in OP50, *n* = 62; *dmsr-7* in OP50, *n* = 62. For experiments in **B** and **C** graphs are mean + s.e.m. For experiments in **D** and **E**, box plots’ center line denotes median; box range denotes 25–75th percentiles; whiskers denote minimum–maximum values. *P < 0.05, **P < 0.01, ***P < 0.001, ns, not significant, (**B**, **C**) by one-way ANOVA with Dunnett’s post-hoc test; comparisons to *dmsr-7(ky1067)*, or (**D**, **E**) by two-way ANOVA with Tukey’s post-hoc test.

The RIM and RIC interneurons produce the neurotransmitters glutamate, tyramine and octopamine, as well as several neuropeptides (Alkema et al., 2005; Serrano-Saiz et al., 2013; Taylor et al., 2021). A loss-of-function mutation in the tyrosine decarboxylase *tdc-1*, which eliminates both tyramine and octopamine synthesis, fully restored PA14 avoidance in *dmsr-7* mutants, while a *tbh-1* mutation that selectively decreases octopamine partially restored PA14 avoidance (**Figures 5C and S5F)**. As single mutants, both *tdc-1(n3419)* and *tbh-1(n3247)* exhibited normal PA14 avoidance (**Figures 5C and S5F)**.

A transcriptional reporter in which the *tdc-1* promoter drove GFP expression was downregulated in both RIM and RIC after PA14 exposure, consistent with a decrease in tyramine and octopamine signaling (**Figures 5D and 5E**). PA14 did not downregulate *tdc-1*p::GFP in the *dmsr-7(ky1067)* mutant background, suggesting that FLP-1 signaling to DMSR-7 regulates *tdc-1* transcription in RIM and RIC. Our results indicate that antagonism between FLP-1 and tyramine/octopamine neurotransmitters determines the dynamics of PA14 avoidance.

## DISCUSSION

Transitions between behavioral states often engage antagonistic neuromodulators that stabilize alternative circuit states. For example, in the arcuate nucleus of the rodent hypothalamus, antagonistic and interconnected NPY/AgRP and POMC neurons release AgRP and αMSH neuropeptides to direct feeding and satiety, respectively (Kim et al., 2018). In *C. elegans,* antagonism between serotonin and the neuropeptide PDF-1 stabilize opposing dwelling and roaming states, modulating behavior in the timescale of minutes (Flavell et al., 2013). Here we show that PA14 pathogen avoidance is regulated over hours by opposing functions of FLP-1 neuropeptides released by the AVK neurons, and tyramine/octopamine neuromodulators produced by RIM and RIC neurons. This graded behavioral change is associated with enhanced FLP-1 release and changes in transcription affecting both neuromodulators, providing a potential mechanism for slow regulation of behavior.

The activity of the AVK neurons determines the temporal dynamics of PA14 avoidance behavior: manipulations that block AVK synaptic release delay avoidance, and manipulations that enhance AVK synaptic release accelerate avoidance. The high potency receptor DMSR-7 in RIM and RIC neurons is one target of FLP-1 peptides, but there are likely additional receptors and neurons that mediate FLP-1 effects on pathogen avoidance. DMSR-7 is related to inhibitory Gi/Go-coupled receptors, which are best known in *C. elegans* for their negative effects on synaptic release (Nurrish et al., 1999; Tanis et al., 2008), but can have other effects, including regulation of CREB transcription factors (Suo et al., 2009). Further experiments are required to determine the cell biological targets of DMSR-7 in RIM and RIC. FLP-1 peptides from AVK can also modulate body posture and locomotion in the timescale of seconds (Hums et al., 2016; Oranth et al., 2018), through the NPR-6 and FRPR-7 receptors, which do not affect pathogen avoidance. Other receptors for FLP-1 neuropeptides remain to be characterized.

We suggest that AVK neurons conduct damage or physiological signals associated with infection from the intestine to the brain (Melo and Ruvkun, 2012; Vasquez-Rifo et al., 2020), thereby linking pathogenesis with the neuronal circuits that modify subsequent responses to PA14. The cell bodies of the AVK neurons are located at the junction of the pharynx and the intestine, and their axons run through the nerve ring, where they contact RIM and RIC, and then down the body alongside the intestine, where they are well-placed to detect changes in intestinal physiology induced by PA14 infection. The critical intestinal signal(s) induced by infection remain unknown, but have been hypothesized to include physical distension, altered intestinal gene expression, and direct physical damage (Dunbar et al., 2012; Fletcher et al., 2019; Melo and Ruvkun, 2012; Singh and Aballay, 2019).

RIM and RIC neurons integrate multiple signals related to pathogen avoidance. In addition to detecting FLP-1 neuropeptides, they function as relays for external PCN and pyochelin signals sensed by ASJ neurons and propagated by DAF-7/TGF-β signaling to RIM/RIC, where the transcriptional regulator DAF-3 acts on unknown targets to alter oxygen responses (Meisel et al., 2014). Despite convergence on RIM and RIC, *daf-7* and *flp-1* have genetically distinct functions: *daf-7* defects are suppressed by mutations in *daf-3,* but not by *tdc-1* mutations that disrupt octopamine and tyramine synthesis (Meisel et al., 2014), whereas *flp-1* and *dmsr-7* defects are suppressed by *tdc-1* but not by *daf-3.* Thus neuropeptide (FLP-1) and cytokine (TGF-β/DAF-7) pathways for sickness behavior affect RIC and RIM differently. Interestingly, suppression of octopaminergic signaling from RIC neurons can enhance innate immune responses against PA14 infection, suggesting that reverse signaling between the nervous system and somatic tissues may contribute to protection from the pathogen (Sellegounder et al., 2018).

Ascertaining bacterial food quality is an ecologically relevant challenge for *C. elegans*. A fifth of the bacteria co-isolated with *C. elegans* from its natural habitat are from the genus *Pseudomonas,* including both high-quality nutritious strains and pathogenic strains that kill the animal (Samuel et al., 2016). *C. elegans* distinguishes between these bacteria by learned behaviors: for example, it is initially attracted to *Pseudomonas* odors, but consumption of PA14 pathogen generates a negative associative memory that triggers avoidance of both pathogenic and nonpathogenic *Pseudomonas* odors upon subsequent exposure (Zhang et al., 2005). The learned avoidance can be retained over development (Jin et al., 2016) and even transmitted to the animal’s progeny across several generations (Kaletsky et al., 2020; Moore et al., 2019). In an additional example of neuromodulatory remodeling in sickness behavior, learned avoidance of pathogen odor requires the neuromodulator serotonin and is associated with transcriptional upregulation of the serotonin biosynthetic gene *tph-1* (Zhang et al., 2005). The diversity and sophistication of *C. elegans’s* responses to *Pseudomonas* highlight the value of using natural stimuli to probe an animal’s behavioral repertoire (Tinbergen, 1951).

Our results implicate neuromodulator release as a key driver of pathogen avoidance, and the observation that *flp-1, tdc-1,* and *tph-1* are all regulated transcriptionally by infection suggests that changes in neuromodulatory gene expression contribute to pathogen-regulated behaviors. Yet FLP-1, serotonin and tyramine all have acute functions in the regulation of locomotion as well, and the neurons that release them are dynamically regulated on short timescales. Neuropeptide gene expression is regulated by activity in many *C. elegans* neurons and mammalian neurons, suggesting a general role for neuromodulatory transcription in the slow modification of circuit function (Laurent et al., 2015; Rojo Romanos et al., 2017; Uhl and Nishimori, 1990). Future experiments are required to determine the quantitative contributions of gene expression, regulated secretion, and regulated neuronal activity to the antagonism between FLP-1 and tyramine/octopamine in pathogen avoidance.

## METHODS

### Nematode Culture

*C. elegans* strains were maintained at room temperature (20-22°C) on nematode growth medium (NGM) plates seeded with *E. coli* OP50 as food source, as previously described (Brenner, 1974). Wild-type animals were CX0006, a descendant of *C. elegans* Bristol N2. Mutant strains used for the candidate gene screen are described in (**Table S1**). All strains and controls were raised together under identical conditions.

### PA14 cultivation

*P. aeruginosa* PA14 was a gift from J. N. Engel. To obtain single colonies of PA14, 10 cm Luria broth (LB) agar plates were warmed up to 37°C, streaked with a PA14 frozen glycerol stock, and incubated at 37°C for 18-20 hours. The streaked plate was stored at 4°C and used within a week. Single colonies of PA14 were cultured in 3 mL of LB at 25°C for ∼20 hours with agitation (250 rpm). The optical density of the liquid culture was measured at 600 nm (OD_600_). To reduce variability within assays, only liquid cultures in the OD_600_ range 4 – 4.5 were used.

Plates for PA14 avoidance assays (standard plates) were generated by seeding 10 μl of PA14 liquid culture in the center of 6 cm NGM plates, generating a circular bacterial lawn. For full-lawn PA14 plates, 200 uL of PA14 liquid culture was spread to fully cover 6 cm NGM plates and excess liquid was removed with a pipette. Seeded plates were placed on a 25°C incubator for 24 hours and then moved to room temperature for 30-60 minutes before transferring animals. Standard and full-lawn PA14 killing plates were prepared in the same way as described above with the addition of 50 μg/mL 5-fluorodeoxyuridine (FUdR) to the NGM to sterilize the animals.

### PA14 avoidance assay

Avoidance assays were initiated by transferring 13-17 animals in the L4 larval stage from their OP50 cultivation plate to a PA14 lawn or a control OP50 lawn. Animals were transferred first to an unseeded NGM plate, allowed to clean themselves from the bacteria for two minutes, then transferred to the assay plate next to the bacterial lawn. Assay plates were placed in a temperature-controlled behavioral setup (21°C) inside a behavioral room (21°C). The L4 to adult molt occurred approximately four hours after transfer, implying that avoidance behavior was detected in young adult animals. Four 15 MP cameras (PL-D7715, Pixelink) were used to record up to 4 assay plates per camera at a rate of 1 frame per minute for 20 hours. After 20 hours, the number of animals on each plate was counted and plates where 3 or more animals were missing were discarded. Custom made MATLAB code (deposited into GitHub) was used to quantify the Avoidance Ratio, the number of animals outside the bacterial lawn over the number of animals at the end of the assays, every 30 minutes.

Although reproducible at a qualitative level, the avoidance assays showed some quantitative variability from month to month, especially at intermediate timepoints. To control for this issue, mutant and transgenic strains were compared to matched controls collected at the same time in all figures and statistical analyses.

### PA14 killing assay

For standard PA14 killing assays, 17-20 animals of each genotype were assayed in triplicate following transfer to a PA14 lawn as described above. For full-lawn PA14 killing assays, 45-60 animals of each genotype were assayed in duplicate. Plates were placed on a 25°C incubator and survivors scored at least twice a day.

### *egl-3* floxed allele and cell-specific inactivation

CRISPR/Cas9 was used for replacing the *egl-3* locus with a donor template by homology directed repair. The donor template contained, in the following order, a left homology arm, a LoxP site 70 base pairs upstream of the *egl-3* start codon, the endogenous *egl-3* gene and its 3’ UTR, a loxP site 852 base pairs downstream of the *egl-3* stop codon, mCherry sequence and the *tbb-2* 3’ UTR, the *rps-0* promoter to drive expression of the hygromycin resistance (hygR) gene and the *unc-54* 3’UTR, and a right homology arm. The map of the plasmid that contains the repair template can be found in the following link (https://benchling.com/s/seq-mO0FvEuUCROr1JZ8ikIN/edit).

Cre expression was accomplished using cell-selective transgenes. The expression patterns of *flp-*1p(1.5 kb)::Cre (*n* = 15), flp*-1*p(513 bp)::Cre (*n* = 26), and *twk-47*p::Cre (*n* = 19) were characterized using a pan-neuronal Cre-reporter strain (h20p::loxP::NLS::mCherry::loxP::NLS::GFP), where Cre expression in a given neuron at any time in development exchanges NLS-mCherry expression for NLS-GFP. We found that *flp-1*p(1.5 kb)::Cre, *flp-1*p(513 bp)::Cre, and *twk-47*p::Cre were expressed only in AVK in 8/15, 24/26, and 19/19 animals, respectively. The interneurons SMBD and SMDD were inconsistently labeled by *flp-1*p(1.5 kb)::Cre and *flp-1*p(513 bp)::Cre. GFP reporters made with *flp*-*1*p(1.5 kb) or *flp-1*p(513 bp) were only expressed in AVK in larval and adult stages as shown in **Figures 3C and S3C**, and also shown by others (Jia and Sieburth, 2021; Li et al., 1999; Oranth et al., 2018).

In the *egl-3* floxed strain, inactivation of *egl-3* in AVK using *flp-1*p(513 bp)::Cre and *twk-47*p::Cre was confirmed by detection of the mCherry reporter.

### Generation of transgenic strains

Transgenic strains, including rescue strains and Cre-expressing strains, were generated following standard microinjection protocols (Mello and Fire, 1995; Mello et al., 1991). A mix of genomic DNA or cDNA clones, a co-injection marker (*elt-2*p::nls-GFP, *myo-3*p::mCherry and/or *myo-2*p::mCherry), and empty pSM vector was injected into adult hermaphrodites. Two to four independent transgenic strains were isolated and characterized to account for variability in transgene expression.

### CRISPR/Cas9-generated mutant strains

Putative null mutants were generated using CRISPR/Cas9 to insert a stop knock-in (STOP-IN) cassette that creates frameshifts and stop codons in all reading frames of the target gene (Wang et al., 2018). Below, the sequences of the insertions, with 20 bp flanks, shown on (+) strand (Insertions are underlined and the sequence containing the stop codons in bold).

*dmsr-1(ky1053)* mutation: GTATATAATTTCTACCATCC<u>GGGAAGTTTGTCCAGAGCAGAGG**TGACTAAGTGATAA**GCTA GC</u>TATACATGCCTACTTATCAA

*dmsr-5(ky1059)* mutation: GCAGGGTCTACACAGTATTG<u>GGGAAGTTTGTCCAGAGCAGAGG**TGACTAAGTGATAA**GCT AGC</u>CATCGGTATTTATGTCTTTT

*dmsr-6(ky1065)* mutation: TTCTTGTCATTCATCCTATG<u>GGGAAGTTTGTCCAGAGCAGAGG**TGACTAAGTGATAA**GCTA GC</u>TGTTCTTGGATTAGCTGCCA

*dmsr-7(ky1067)* mutation: TATCTTCACCAATTTTGTGC<u>GGGAAGTTTGTCCAGAGCAGAGGTGAGGGAAGTTTGTCCAG AGCAGAGG**TGACTAAGTGATAA**GCTAGC</u>ATGTGGCAGTTTTATCGAGA

### Intersectional Cre/Lox rescue

Cell-specific rescues with transgenes were generated with an intersectional strategy using the Cre-Lox system to restrict *dmsr-7* expression to a subset of its endogenous expression pattern. Gibson Assembly (New England Biolabs) was used to subclone fragments of the pSM-inv[SL2-GFP] vector (Flavell et al., 2013) to generate an inactive, inverted *dmsr-7* construct (**Figure 4C**). First, a genomic fragment of *dmsr-7* including 5033 bp of the upstream promoter region and 815 bp of the open reading frame was cloned upstream of the first Lox2272 and LoxP sites. Then, the remaining *dmsr-7* fragment was cloned in an inverted orientation between the inverted SL2-GFP and the second Lox2272 and inverted LoxP.

Transgenic *dmsr-7(ky1067)* mutants expressing the inverted *dmsr-7* construct were injected with plasmids expressing Cre under cell-specific promoters. In these transgenic lines, the *dmsr-7* genomic fragment is reconstituted only in the cells that express both Cre and the inverted *dmsr-7* construct. Reconstitution of the *dmsr-7* rescue fragment was confirmed by GFP expression.

### Histamine silencing experiments

A stock solution of 1 M histamine-dihydrochloride (Sigma-Aldrich) in deionized water was filtered and stored at −20°C. NGM-Histamine plates were prepared by adding 1 M histamine to NGM agar to a final concentration of 10 mM. Plates were left at room temperature for a day and stored at 4°C for up to a month. All animals expressing HisCl1 were assayed in plates with and without histamine alongside the corresponding parental strain that do not express the transgene.

### GPCR activation assays

GPCR activation assays were performed using CHO-K1 cells stably expressing the luminescent Ca^2+^ indicator aequorin and the promiscuous Gα16 protein (ES-000-A24 cell line, PerkinElmer). The cells were transfected with a *dmsr-7*/pcDNA3.1 plasmid using lipofectamine LTX and Plus reagent (Invitrogen) and grown overnight at 37°C. One day post-transfection, they were supplemented with fresh culture medium and shifted to 28°C overnight. Two days post-transfection, cells were collected in bovine serum albumin (BSA) medium (DMEM/F12 without phenol red, supplemented with 0.1% BSA) at a density of 5 million cells per mL, and loaded with 5 µM coelenterazine h (Invitrogen) for 4 hours at room temperature. After a 10-fold dilution, cells (25,000/well) were challenged with synthetic peptides reconstituted in DMEM/BSA medium and luminescence was measured on a MicroBeta LumiJet luminometer (PerkinElmer). Luminescence was measured for 30 s at a wavelength of 469 nm. After 30 s, 0.1% Triton X-100 (Merck) was added to lyse the cells, resulting in a maximal Ca^2+^ response that was measured for 30 s. Concentration-response measurements were done in triplicate on two independent days. For each peptide concentration, a relative calcium response (%) compared to the maximum peptide-evoked response (100% activation) was calculated. Concentration-response data were then fitted in function of log[peptide]. EC_50_ values were calculated from concentration-response curves in GraphPad Prism 7 by fitting a 3- or 4-parameter concentration-response curve.

### Imaging and fluorescence quantification

Transgenic animals expressing *flp-1*p(1.5 kb)::GFP, *flp-1*p(513 bp)::FLP-1(cDNA)::mCherry, or *tdc-1*p::GFP were placed into full-lawn PA14 plates to eliminate variations due to avoidance behavior. Transgenic animals grown on *E. coli* OP50 plates were randomly transferred to a full-lawn PA14 plate or a control OP50 plate, and incubated for 4 or 10 hours at 20°C before imaging.

Live animals were mounted on 2% agarose pads containing 5 mM sodium azide. Fluorescence was collected with a 40X objective on a Zeiss Axio Imager.Z1 Apotome microscope with a Zeiss AxioCam MRm CCD camera (**Figures 3E and 3F**), or a Zeiss Inverted Axio Observer Z1 LSM 780 laser scanning confocal microscope with a 40x objective (**Figures 3C and 3D**). Imaging settings were constant across all experiments. Images were processed using ImageJ (NIH, Bethesda, USA) to generate a maximum-intensity Z-projection. Quantification of fluorescence levels was performed by drawing a region of interest (ROI) around the cell body and measuring mean gray scale values.

### Statistical Methods

Statistical analyzes were performed using GraphPad Prism. For the candidate gene screen, sample size was calculated to test if, at the 20-hour timepoint, the mean of the wildtype control and any of up to five mutant strains tested simultaneously were equal. By using preliminary data collected with wildtype animals (mean: 0.9253, std: 0.0672, *n* = 12), we determined that all mutant strains should be tested at least 6 times in order to find significant differences at least 2 standard deviations below the mean of wildtype controls. A shortcoming of this calculation is the assumption that the mutant strains would have a standard deviation equal to that of the wildtype group. The online calculator can be found on the following link (http://powerandsamplesize.com/Calculators/Compare-k-Means/1-Way-ANOVA-Pairwise).

## ACKNOWLEDGMENTS

We thank Margaret Ebert, Likui Feng, James Lee, and Christine Li for comments on this manuscript. This work was supported by a grant from the Chan Zuckerberg Initiative to C.I.B., grants from the Research Foundation Flanders (G093419N) and KU Leuven Research Council (Grant C19/19/003) to I.B, and grants from NIH (R01NS076558 and DP1NS111778) and a HHMI Scholar Award to D.A.C.R.

## CONTRIBUTIONS

J.M.S. and C.I.B. designed experiments. J.M.S conducted behavioral, genetic, and molecular experiments. J.H. and D.A.C.R. generated the *egl-3* floxed strain. E.V. and I.B. performed *in vitro* experiments to identify FLP-1 receptors. J.M.S. and C.I.B. analyzed and interpreted results. J.M.S. and C.I.B. wrote the paper with input from all authors.

## SUPPLEMENTAL FIGURE AND TABLE LEGENDS

**Figure S1.**
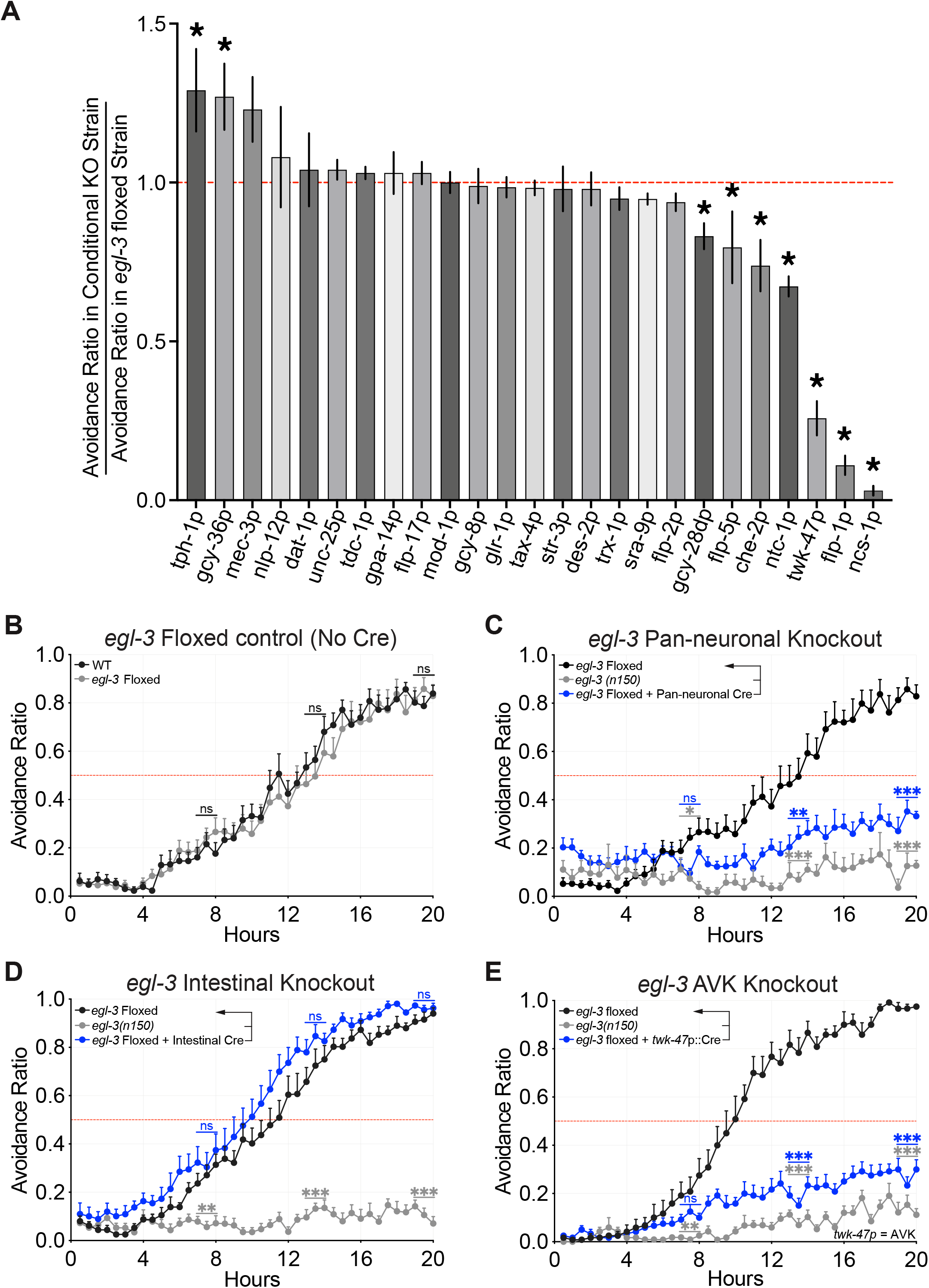
Cell-specific inactivation of *egl-3*. **A**, Cell-specific *egl-3* knockout screen for identifying neurons involved in PA14 avoidance. For each group, the Avoidance Ratio was calculated 20 hours after exposure and normalized to *egl-3*-Floxed controls. For detailed results, see **Table S2**. **B**, *egl-3*-Floxed animals exhibit wild-type PA14 avoidance. **C**, Pan-neuronal *egl-3* knockout causes PA14 avoidance defects. **D**, Intestinal *egl-3* knockout has no effect in PA14 avoidance. **E**, AVK *egl-3* knockout causes PA14 avoidance defects, resembling *egl-3(n150)* mutants. For experiments in **B**, *n* = 9 assays for all groups; for experiments in **D**, **E**, *n* = 8 assays for all groups; for experiments in **C**, *n* = 9 assays for *egl-3*-floxed, *n* = 4 assays for *egl-3(n150)*, *n* = 8 assays for *egl-3*-floxed + Pan-neuronal::Cre. Graphs are mean + s.e.m. *P < 0.05, **P < 0.01, ***P < 0.001, ns, not significant, (**A**, **C**-**E**) by one-way ANOVA with Dunnett’s post-hoc test; comparisons to *egl-3* floxed, or (**A**, **B**) by unpaired two-tailed *t*-test.

**Figure S2.**
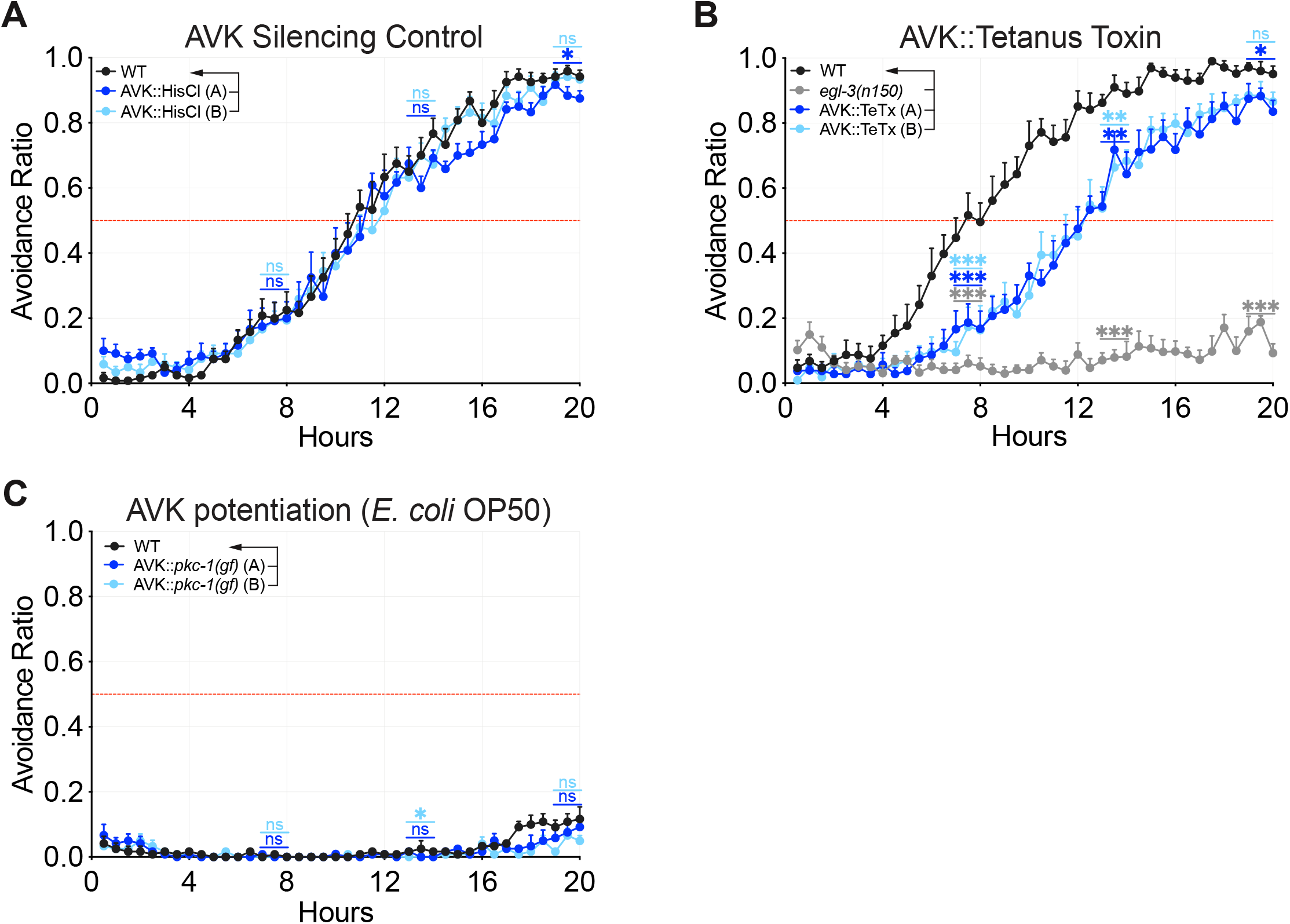
Manipulations of AVK neurons and controls. **A**, Animals expressing HisCl1 in the AVK neurons behave like wild-type controls in plates without histamine. **B**, Expression of the tetanus toxin light chain in AVK neurons delays PA14 avoidance. **C**, Potentiation of dense core vesicle release by expression of *pkc-1(gf)* in the AVK neurons does not induce avoidance to non-pathogenic OP50 lawns. For experiments in **A** and **C**, *n* = 8 assays for all groups; for experiments in **B**, *n* = 7 assays for all groups. Graphs are mean + s.e.m. *P < 0.05, **P < 0.01, ***P < 0.001, ns, not significant, by one-way ANOVA with Dunnett’s post-hoc test; comparisons to WT.

**Figure S3.**
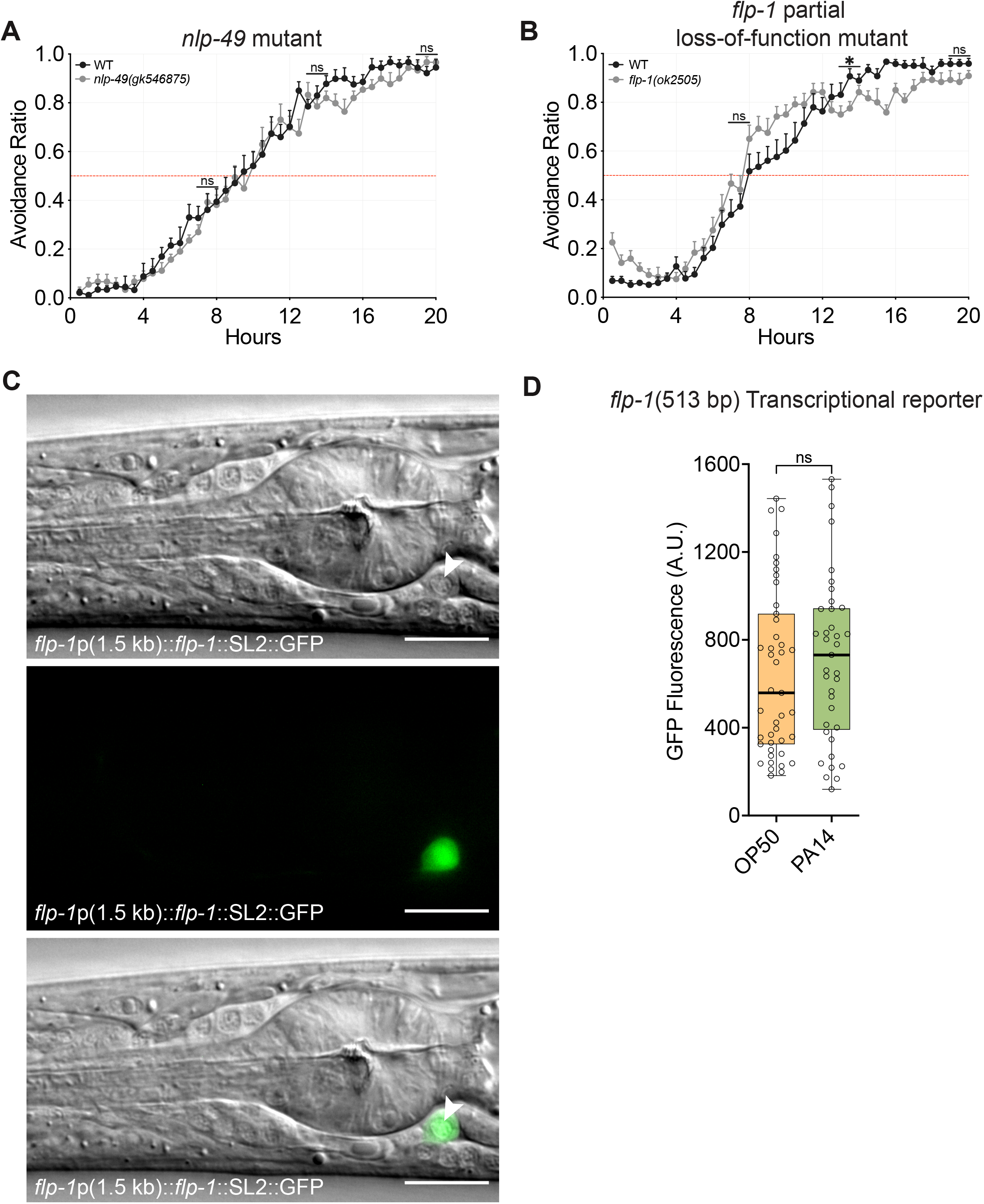
n*l*p*-49* and *flp-1* mutants, and controls. **A**, *nlp-49(gk546875)* loss-of-function mutants have wild-type PA14 avoidance. **B**, *flp-1(ok2505)* partial loss-of-function mutants lack FLP-1 RYamide peptides: FLP-1-8 (KPNFMRY) and FLP-1-9 (PNFMRY). These mutants have wild-type PA14 avoidance, suggesting these peptides are not necessary for the behavior. **C**, The *flp-1*p(1.5 kb)::*flp-1*::SL2::GFP rescue construct is only expressed in the AVK neurons. Top panel, Nomarski image, white arrowhead shows the cell body of AVK left; Medium panel, Fluorescence image (GFP); Bottom panel, overlaid images. Scale bar 10 um. **D**, The *flp-1*p(513 bp)::GFP transcriptional reporter was not significantly induced 4 hours after exposure to PA14, although there was a trend toward increased expression. **E**, For experiments in **A** and **B**, *n* = 8 assays for all groups; for experiments in **D**, *n* = 43 for OP50 group, *n* = 37 for PA14 group. Graphs are mean + s.e.m. *P < 0.05, ns, not significant, (**A**, **B**) by one-way ANOVA with Dunnett’s post-hoc test; comparisons to WT, or (**D**) by unpaired two-tailed *t*-test.

**Figure S4.**
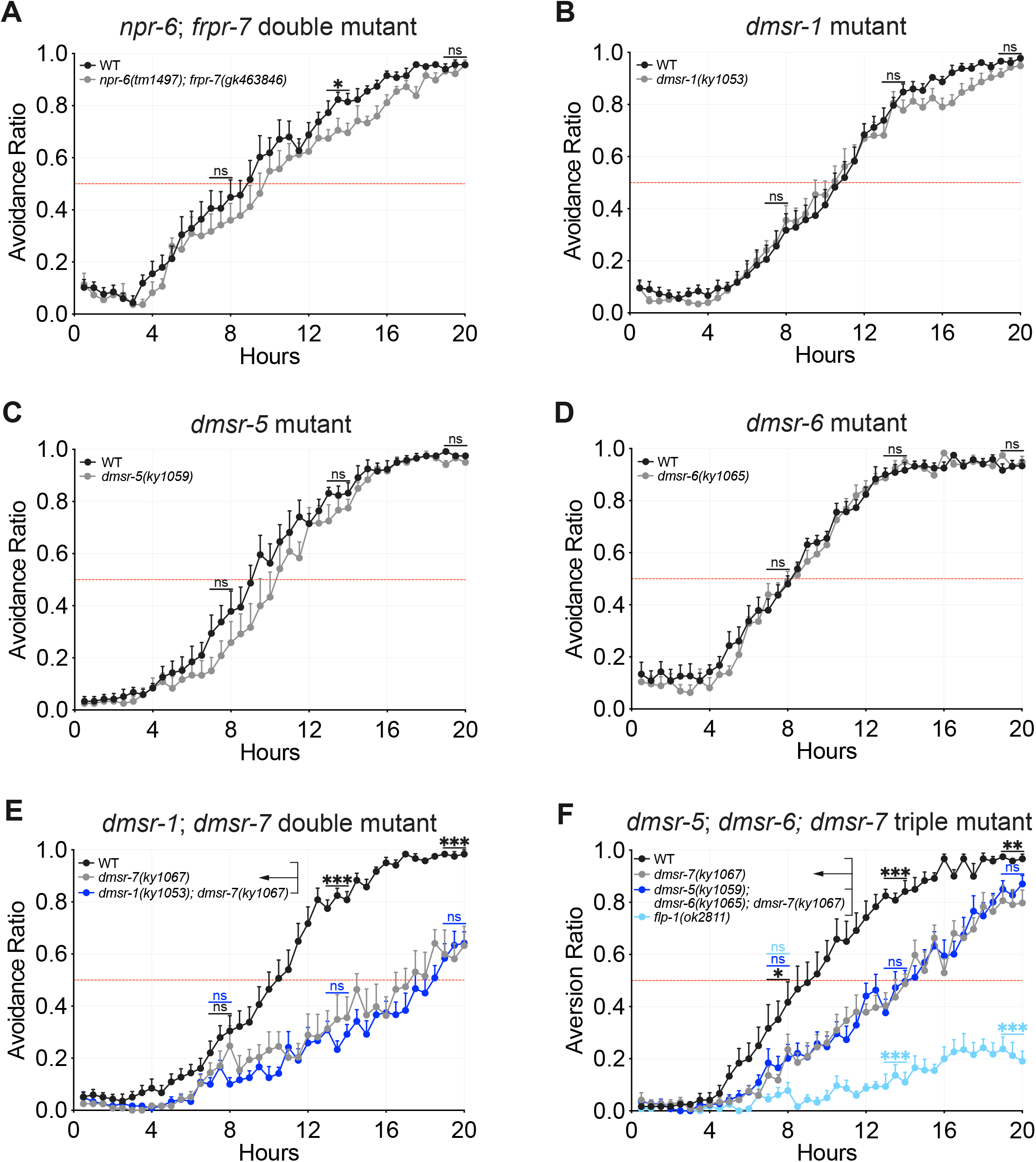
FLP-1 receptor candidates. **A**, *npr-6(tm1497) frpr-7(gk463846)* double mutants affect locomotion parameters such as body bend angles and speed (Oranth et al., 2018), but exhibit normal PA14 avoidance. **B**-**D**, *dmsr-1(ky1053)* (**B**), *dmsr-5(ky1059)* (**C**), and *dmsr-6(ky1065)* (**D**) single mutants exhibit normal PA14 avoidance. **E**, **F**, *dmsr-1 dmsr-7* double mutants, and *dmsr-5 dmsr-6 dmsr-7* triple mutants behave like *dmsr-7* single mutants in the PA14 avoidance assay. For experiments in **A**-**F**, *n* = 8 assays for all groups. Graphs are mean + s.e.m. *P < 0.05, **P < 0.01, ***P < 0.001, ns, not significant, (**A**-**D**) by unpaired two-tailed *t*-test, or (**E** by one-way ANOVA with Dunnett’s post-hoc test; comparisons to *dmsr-7(ky1067)*, or (**F**) by one-way ANOVA with Tukey’s post-hoc test; comparisons to *dmsr-7(ky1067)* shown, for all comparisons see **Table S4**.

**Figure S5.**
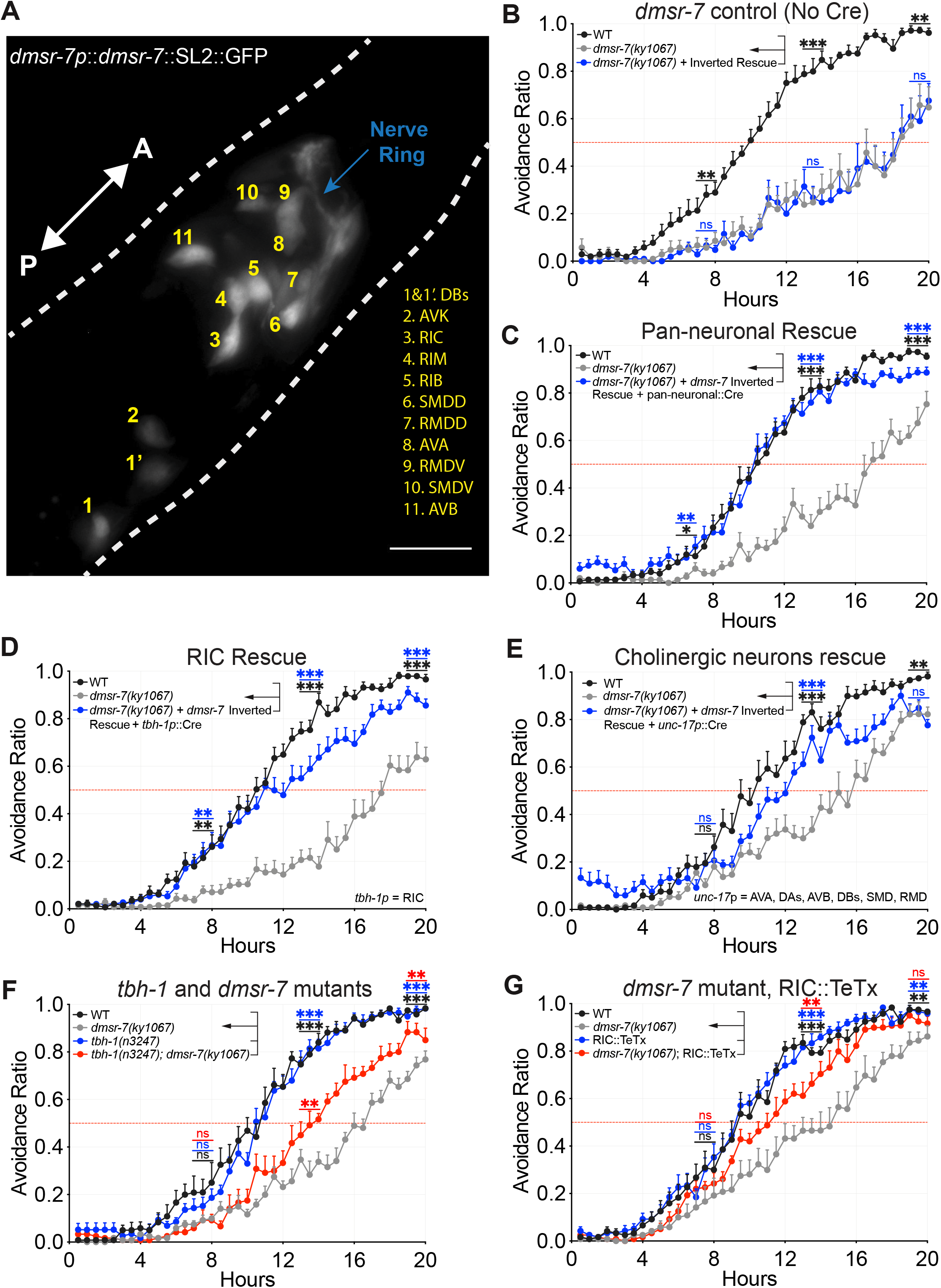
Rescue of *dmsr-7* in subsets of its endogenous expression pattern. **A**, Expression pattern of genomic fragment containing the *dmsr-7* endogenous promoter and the complete *dmsr-7* coding region. GFP was expressed with *dmsr-7* in a bicistronic transcript. Scale bar 10 um. **B**, The *dmsr-7* inverted transgene did not rescue *dmsr-7(ky1067)* PA14 avoidance. **C**, Co-expression of the *dmsr-7* inverted transgene and pan-neuronal Cre rescued *dmsr-7* PA14 avoidance. **D**, **E**, Co-expression of the *dmsr-7* inverted transgene and (**D**) *tbh-1*p::Cre (RIC neurons), or (**E**) *unc-17*p::CRE (AVA, DAs, AVB, DBs, SMD, RMD neurons) resulted in significant rescue of *dmsr-7* PA14 avoidance. **F**, *tbh-1* encodes a tyramine-beta hydroxylase involved in the biosynthesis of octopamine. A mutation in *tbh-1* partially suppresses the *dmsr-7* PA14 avoidance defect. **G**, Expression of tetanus toxin light chain in the RIC interneurons significantly suppresses the *dmsr-7* PA14 avoidance defect. For experiments in **B**, *n* = 7 assays for all groups; for experiments in **C**-**E**, *n* = 10 assays for all groups; for experiments in **F**, **G**, *n* = 8 assays for all groups. Graphs are mean + s.e.m. *P < 0.05, **P < 0.01, ***P < 0.001, ns, not significant, (**B**, **C**) by one-way ANOVA with Dunnett’s post-hoc test; comparisons to *dmsr-7(ky1067)*, or (**D**-**G**) by one-way ANOVA with Tukey’s post-hoc test; comparisons to *dmsr-7(ky1067)* shown, for all comparisons see **Table S4**. Although (**D**, **E**) rescue or (**G**) suppression of the *dmsr-7* mutants attained nominal significance compared to both wild-type and *dmsr-7* controls, we cannot distinguish between variability in transgene expression and biological variability for the partial effect.

**Figure S6.**
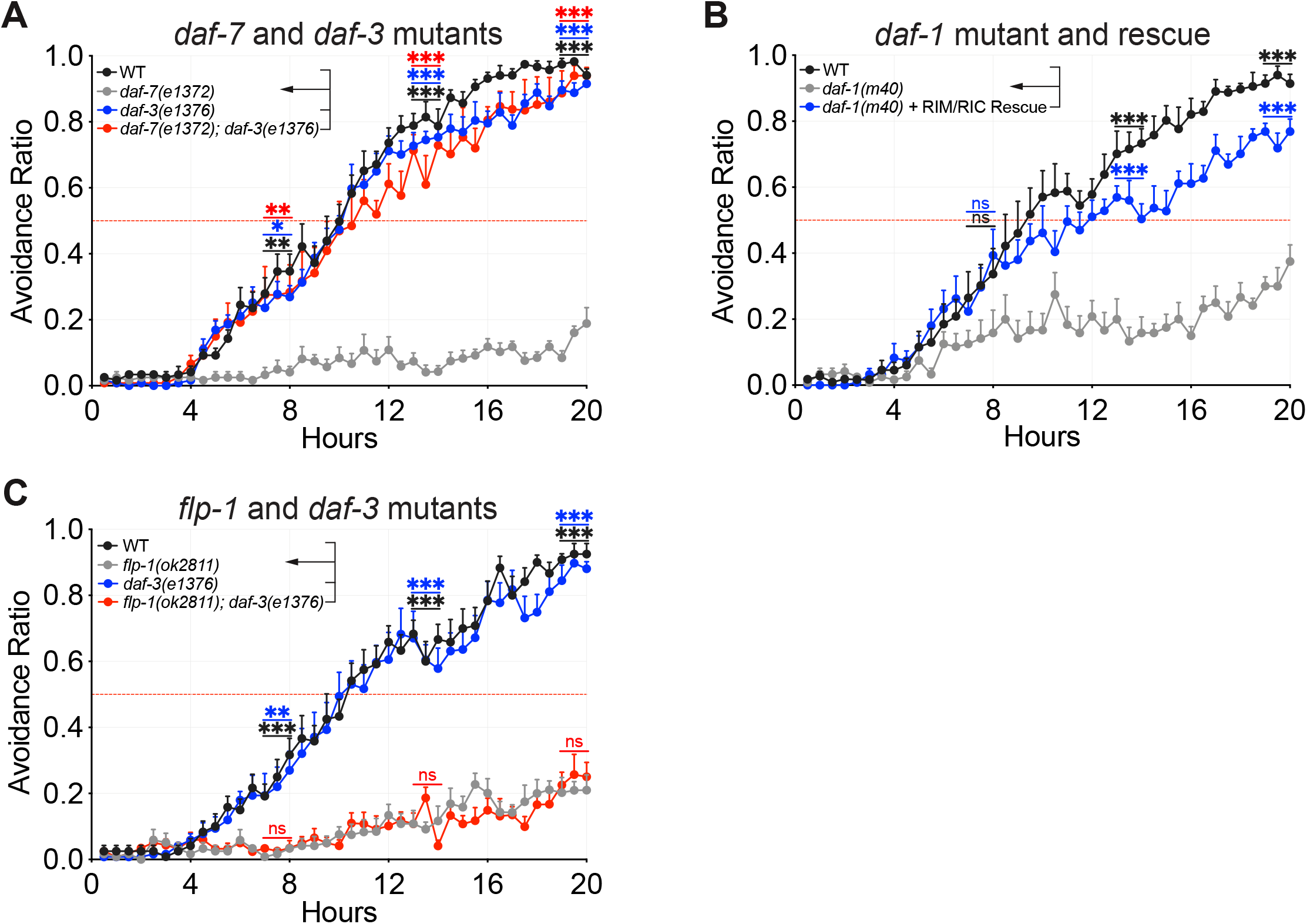
FLP-1 and DAF-7 work through different targets in RIM and RIC. **A**, *daf-3(e1376)* SMAD mutants exhibit wild-type PA14 avoidance and suppress the *daf-7(e1372)* TGF-beta mutant phenotype. **B**, *daf-1(m40)* TGF-beta receptor mutants exhibit a reduction in PA14 avoidance. Rescue of *daf-1* only in the RIM and RIC neurons restores PA14 avoidance to near wild-type levels(Meisel et al., 2014). **C**, A mutation in *daf-3* does not suppress the *flp-1(ok2811)* PA14 avoidance defect. For experiments in **A**-**C**, *n* = 8 assays for all groups. Graphs are mean + s.e.m. *P < 0.05, **P < 0.01, ***P < 0.001, ns, not significant, by one-way ANOVA with Dunnett’s post-hoc test; (**A**) comparisons to *daf-7(e1372)*, (**B**) *daf-1(m40)*, and (**C**) *flp-1(ok2811)*.

**Table S1.**
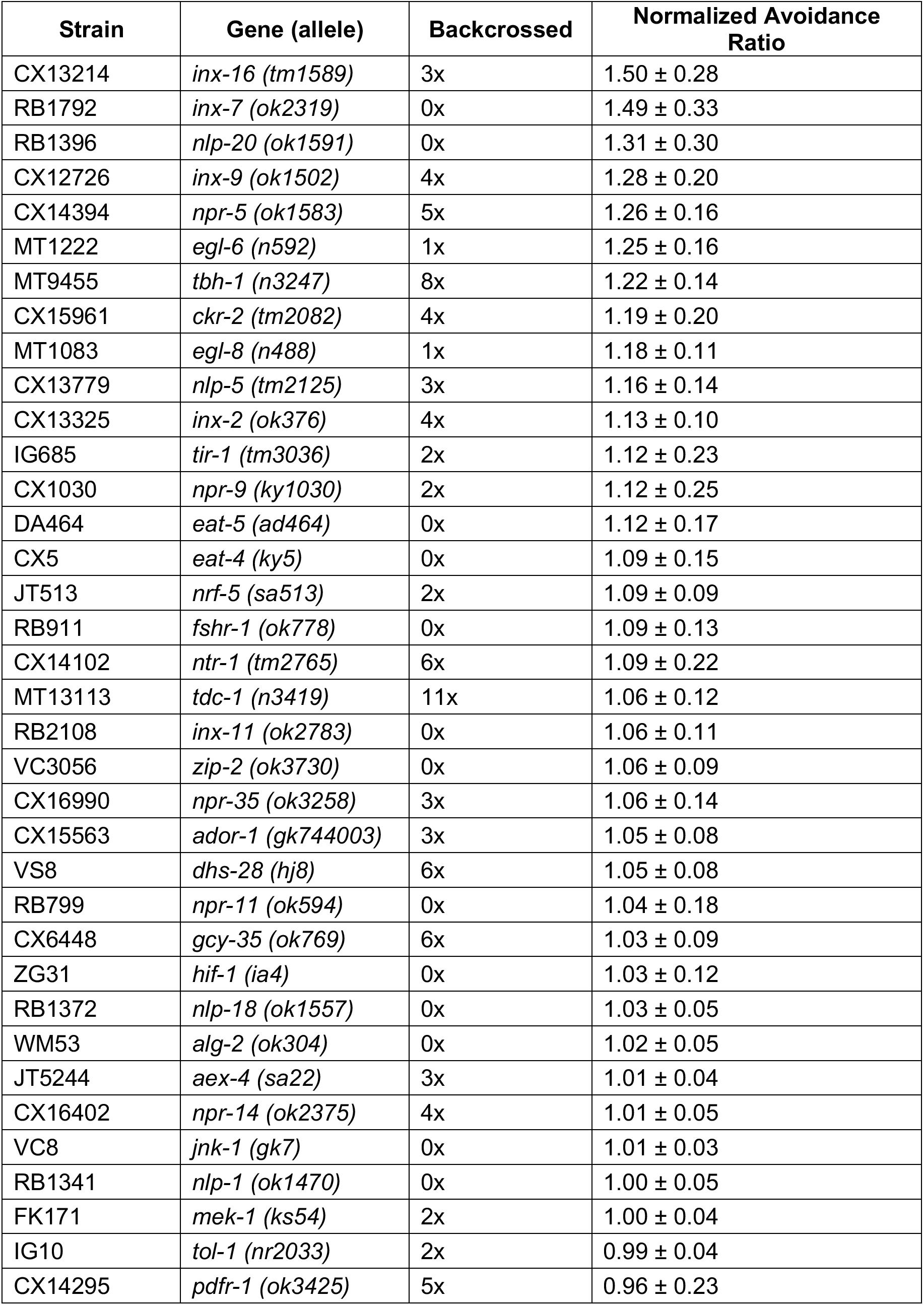
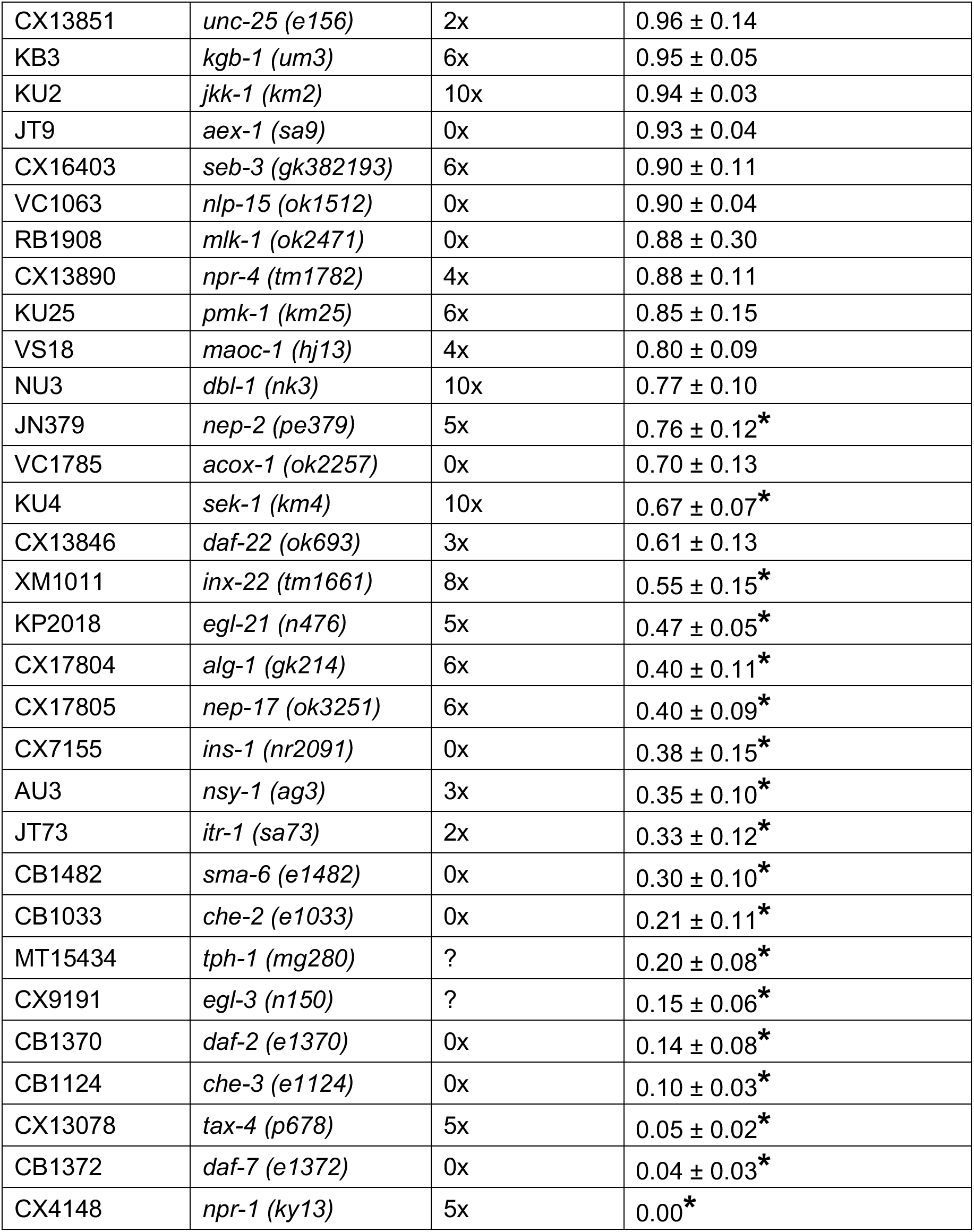
Candidate gene screen for mutants defective in PA14 avoidance. Normalized Avoidance Ratio is the Avoidance Ratio of the mutant strain / Avoidance Ratio of the wild-type control conducted in parallel at 20 hours after exposure to PA14. Note that only one mutant allele was tested per gene, and therefore some phenotypes may not be specific to the gene tested. Data shown is mean ± s.e.m. For all mutants *n* = 6 assays. *P < 0.05 by one-way ANOVA with Dunnett’s post-hoc test; comparisons to respective wild-type controls.

**Table S2.** Unbiased cell-directed neuropeptide screen for neurons required for PA14 avoidance. Normalized Avoidance Ratio is the Avoidance Ratio of the conditional Knockout strain / Avoidance Ratio of *egl-3* floxed controls conducted in parallel at 20 hours after exposure to PA14. Reported expression patterns of relevant promoter fragments were drawn from the literature. Data shown is mean ± s.e.m. For all conditional KO strains, *n* = 6 or 8 assays. *P < 0.05 by one-way ANOVA with Dunnett’s post-hoc test or by unpaired two-tailed *t*-test; comparisons to respective *egl-3* floxed controls.

| Strain name | Promoter driving Cre | Normalized Avoidance Ratio | Neurons targeted |
| --- | --- | --- | --- |
| CX17840 | *tph-1p* | 1.29 ± 0.13* | ADF, NSM, HSN |
| CX17838 | *gcy-36p* | 1.27 ± 0.10* | URX, PQR, AQR |
| CX17836 | *mec-3p* | 1.23 ± 0.10 | ALM, AVM, FLP, PLM, PVD, PVM |
| CX17852 | *nlp-12p* | 1.08 ± 0.16 | DVA |
| CX17830 | *dat-1p* | 1.04 ± 0.12 | ADE, CEPD, CEPV, PDE, OLL |
| CX17858 | *unc-25p* | 1.04 ± 0.03 | RIS, VDs, DDs, AVL, DVB, RME, RIB |
| CX17844 | *tdc-1p* | 1.03 ± 0.02 | RIC, RIM |
| CX17828 | *gpa-14p* | 1.03 ± 0.07 | ADE, ALA, ASH, ASI, ASJ, ASK, AVA, CAN, DVA, PHA, PHB, PVQ, RIA |
| CX17864 | *flp-17p* | 1.03 ± 0.04 | BAG, ASI |
| CX17834 | *mod-1p* | 1.00 ± 0.03 | AIA, AIB, AIY, AIZ, RIC, RID, RIM, RME, DDs, VDs |
| CX17854 | *gcy-8p* | 0.99 ± 0.05 | AFD |
| CX17832 | *glr-1p* | 0.99 ± 0.03 | AIB, AVA, AVB, AVD, AVE, AVG, AVJ, DVC, PVC, PVQ, RIG, RIM, RIS, RMD, RMDD, RMDV, RME, SMDD, SMDV, URY |
| CX17811 | *tax-4p* | 0.98 ± 0.02 | ASJ, AWB, ASI, URX, ASG, AWC, ASK, BAG, ASE, AFD |
| CX17846 | *str-3p* | 0.98 ± 0.07 | ASI |
| CX17924 | *des-2p* | 0.98 ± 0.05 | RID, others |
| CX17842 | *trx-1p* | 0.95 ± 0.04 | ASJ |
| CX17823 | *sra-9p* | 0.95 ± 0.02 | ASK |
| CX17926 | *flp-2p* | 0.94 ± 0.03 | RID, others |
| CX17825 | *gcy-28dp* | 0.83 ± 0.04* | AIA, ASI, AVF |
| CX17860 | *flp-5p* | 0.80 ± 0.11* | RMG, ASE, PVT, I4, M4 neurons |
| CX17819 | *che-2p* | 0.74 ± 0.08* | ASE, ADE, ADF, ADL, AFD, AQR, ASG, ASH, ASI, ASJ, ASK, AWA, AWB, AWC, BAG, CEP, FLP, IL1, IL2, OLL, OLQ, PDE, PHA, PHB, PQR |
| CX17850 | *ntc-1p* | 0.67 ± 0.03* | AFD, AVK, DVA |
| CX18257 | *twk-47p* | 0.26 ± 0.05* | AVK |
| CX18236 | *flp-1(513 bp)p* | 0.11 ± 0.03* | AVK |
| CX17862 | *ncs-1p* | $0.03 \pm 0.01^*$ | ADL, AFD, AVK, ASE, ASG, ASI, AVE, AWB, AWC, BAG, RMG, 1 additional cell anterior to nerve ring, 1 tail neuron |

**Table S3.**
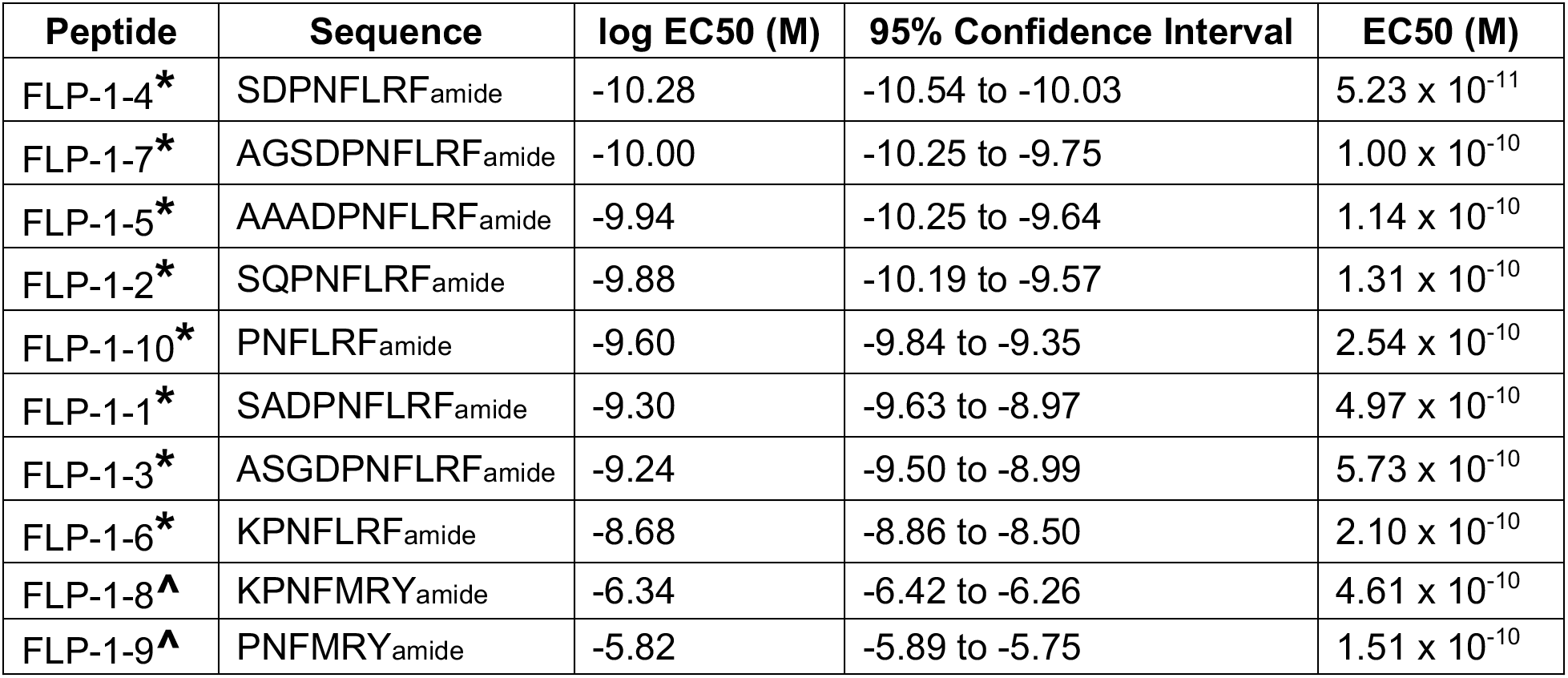
Interactions of DMSR-7 with FLP-1 neuropeptides in CHO-K1 cells. * = FLP-1 RFamide peptide, ^ = FLP-1 RYamide peptide

**Table S4.**
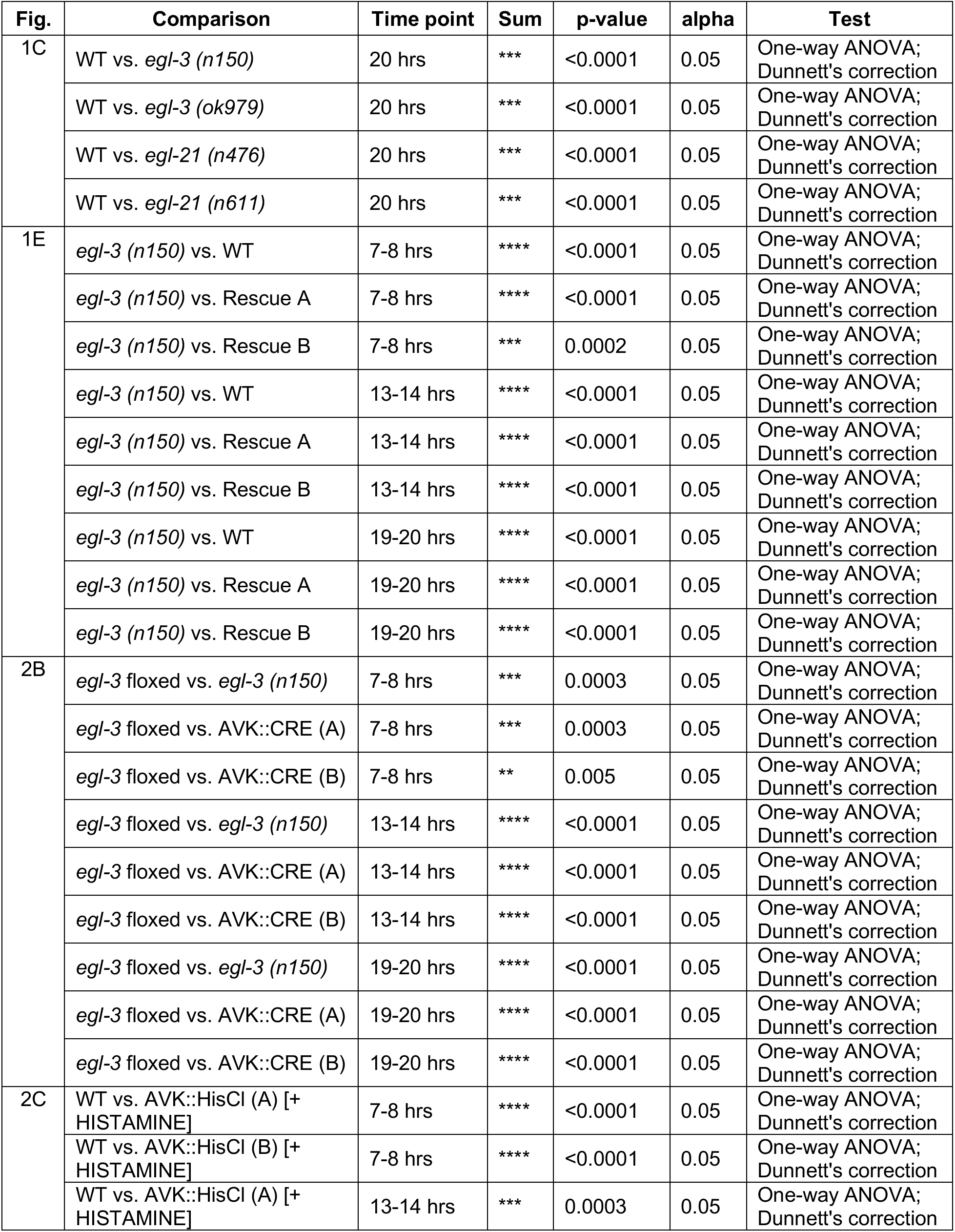
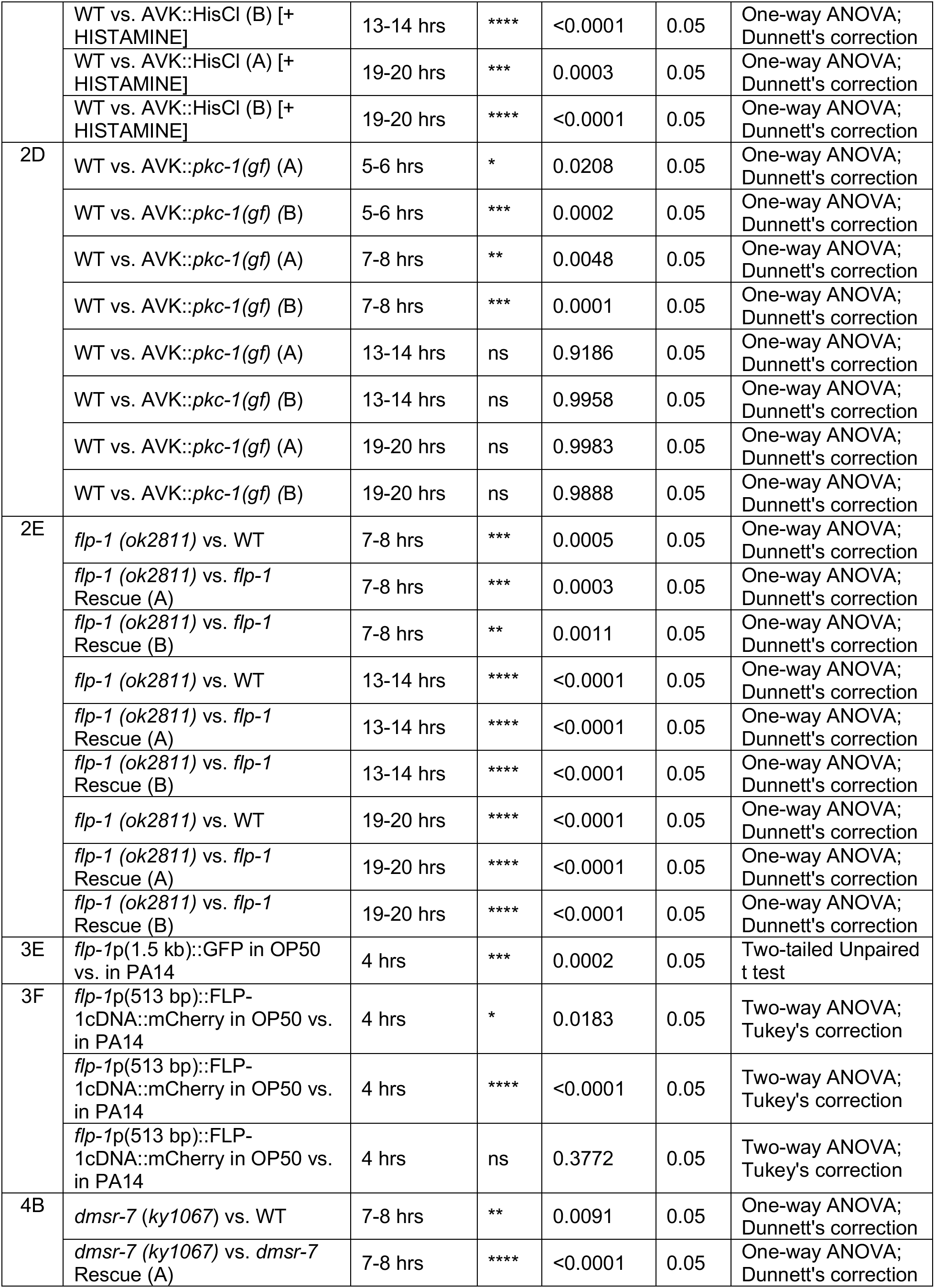
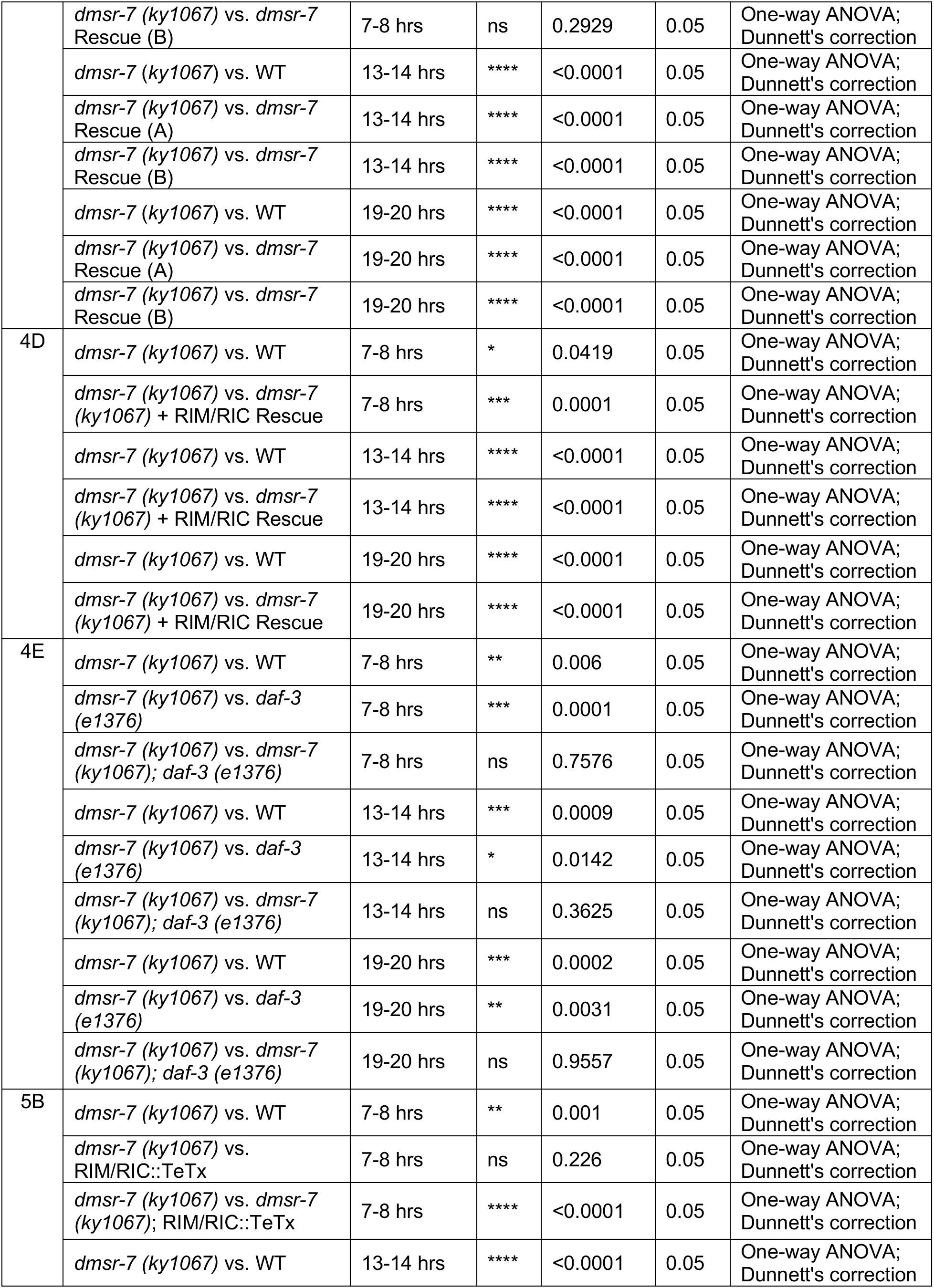
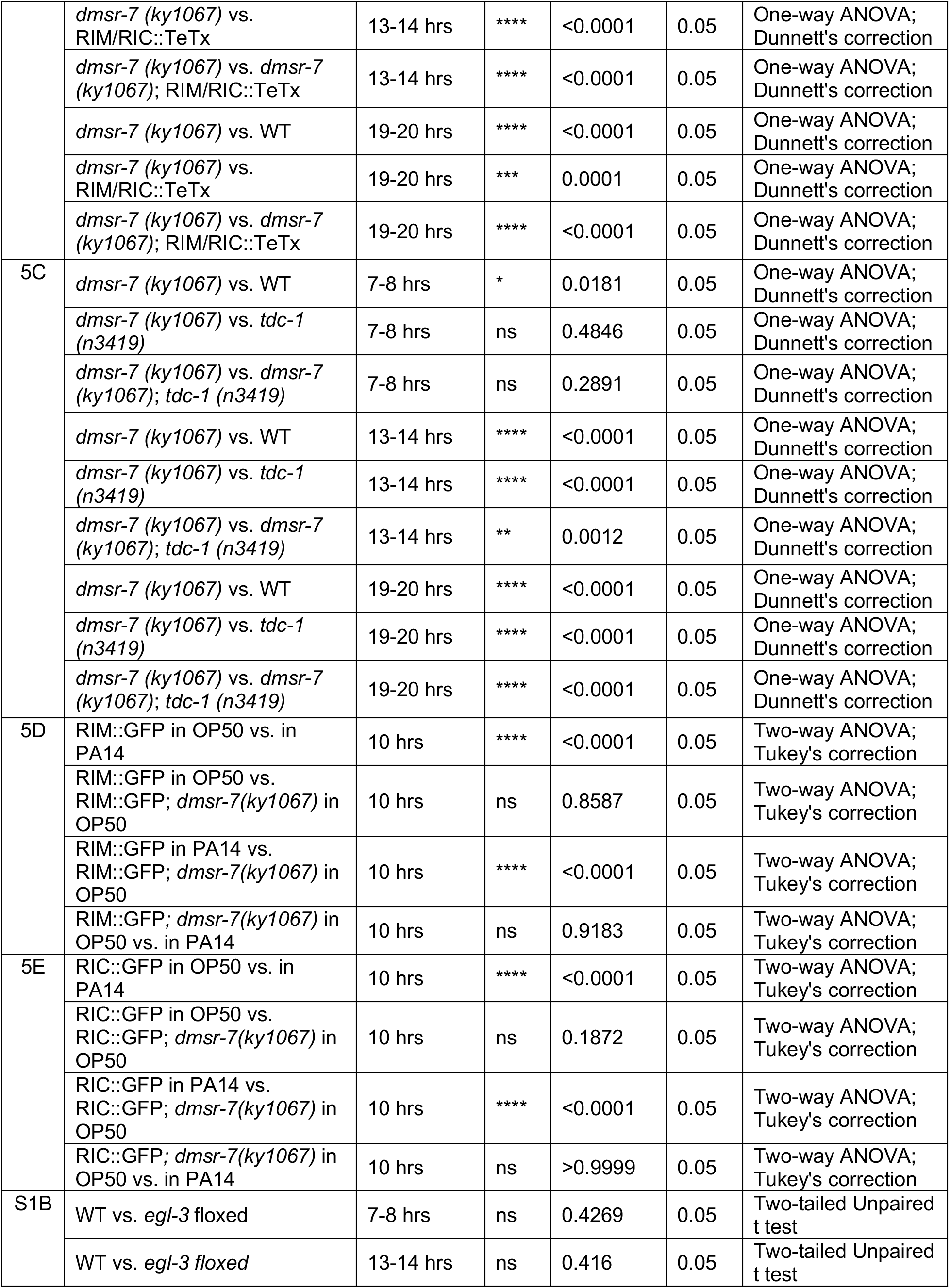
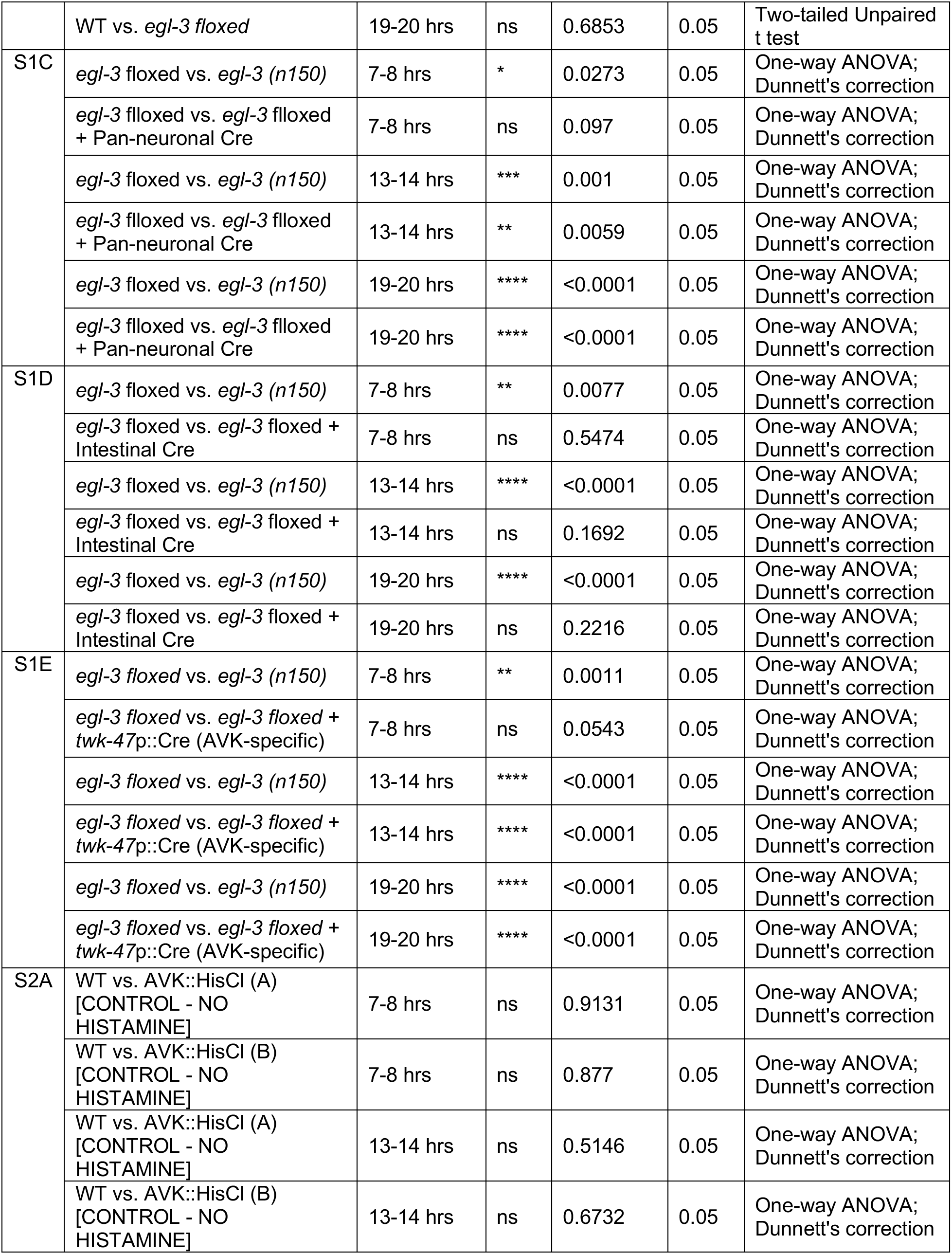
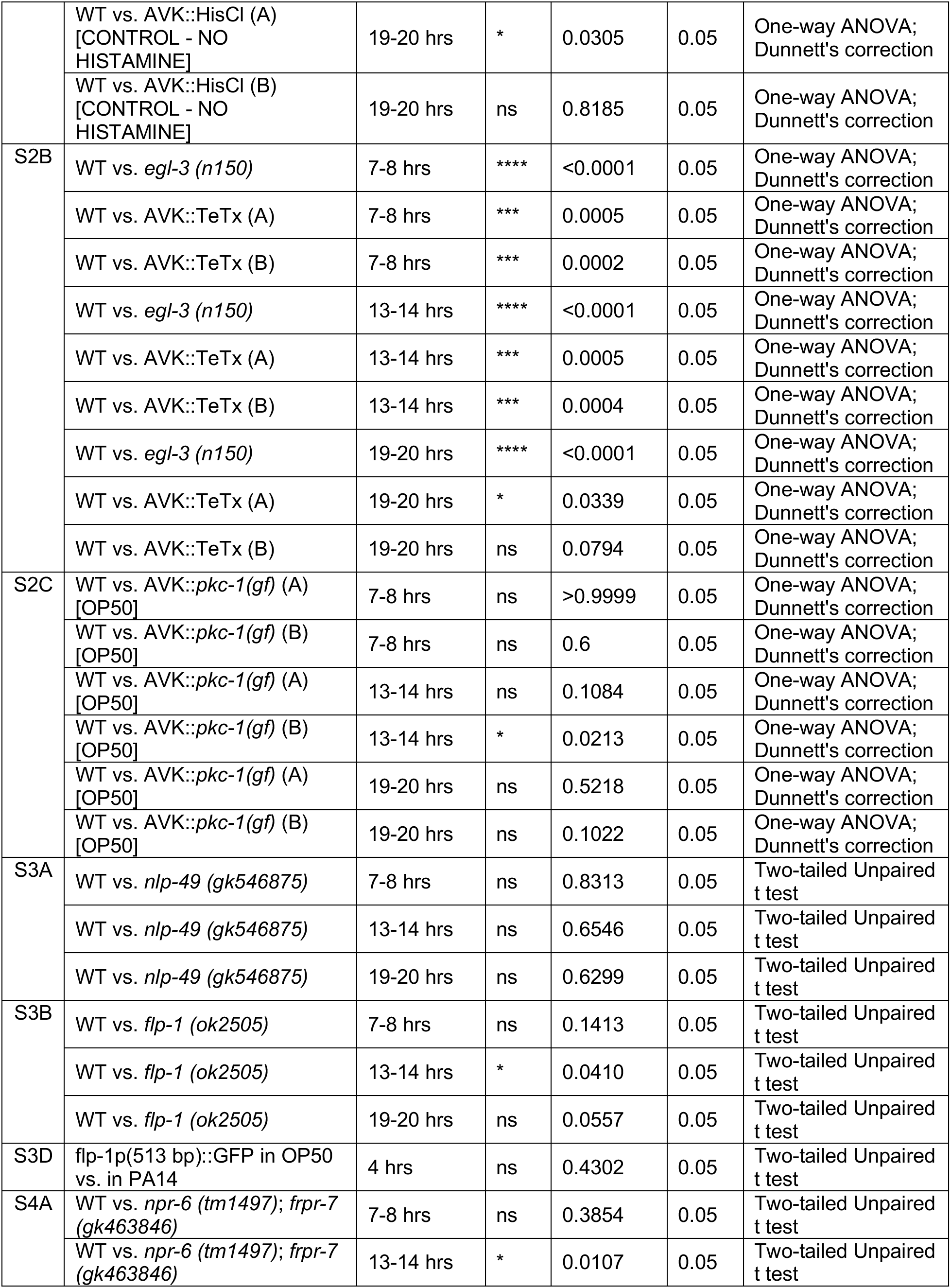
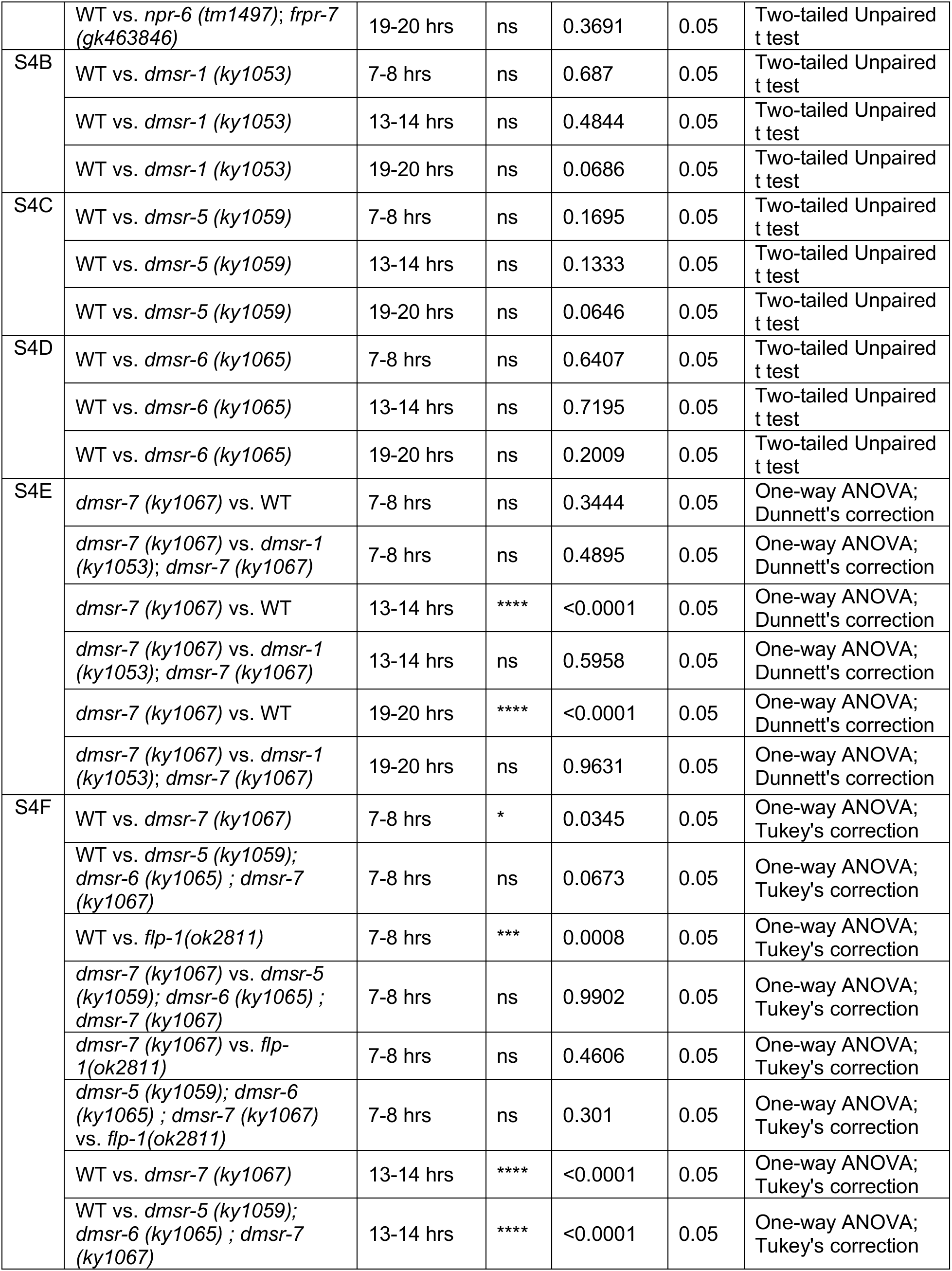
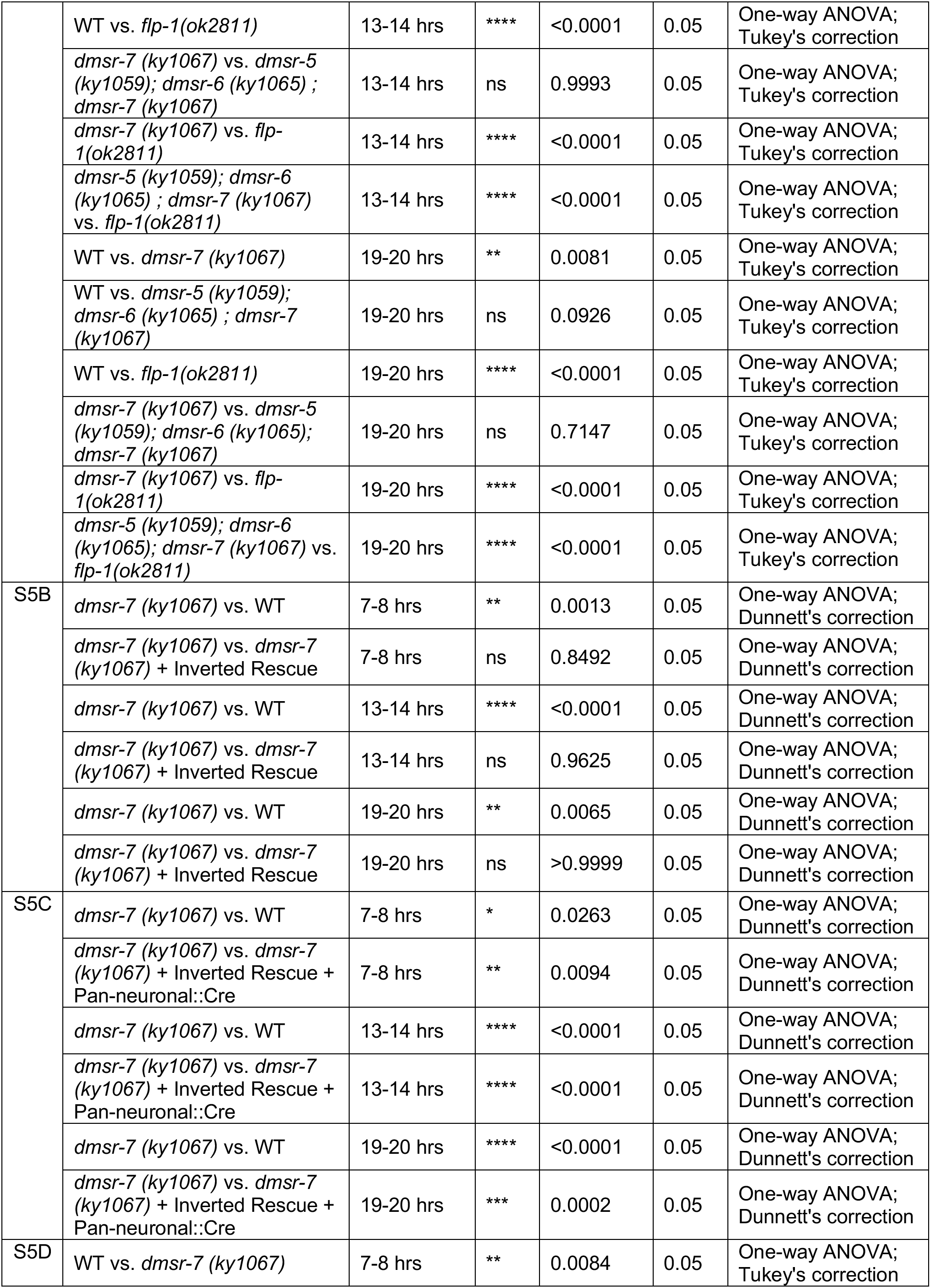
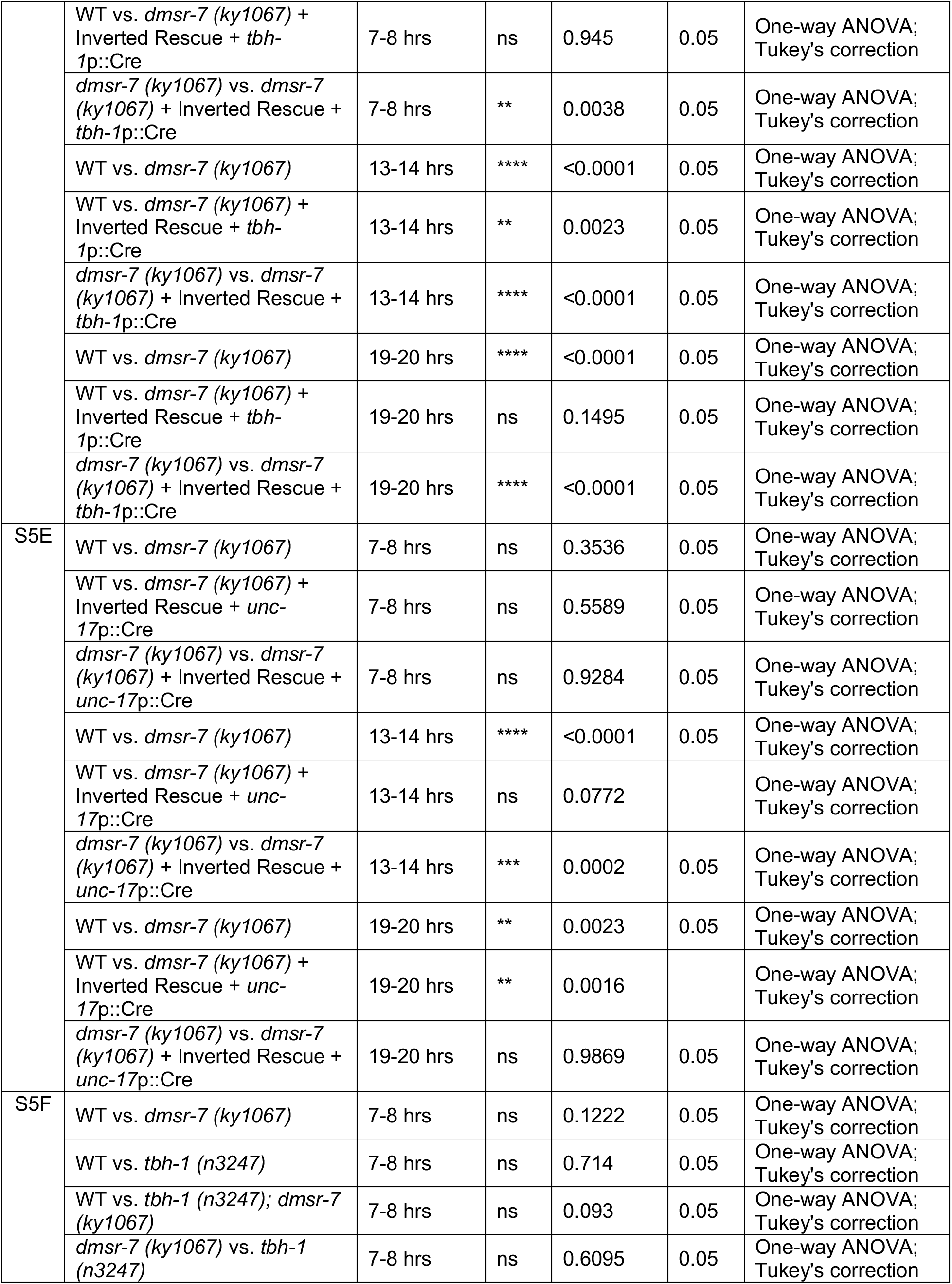
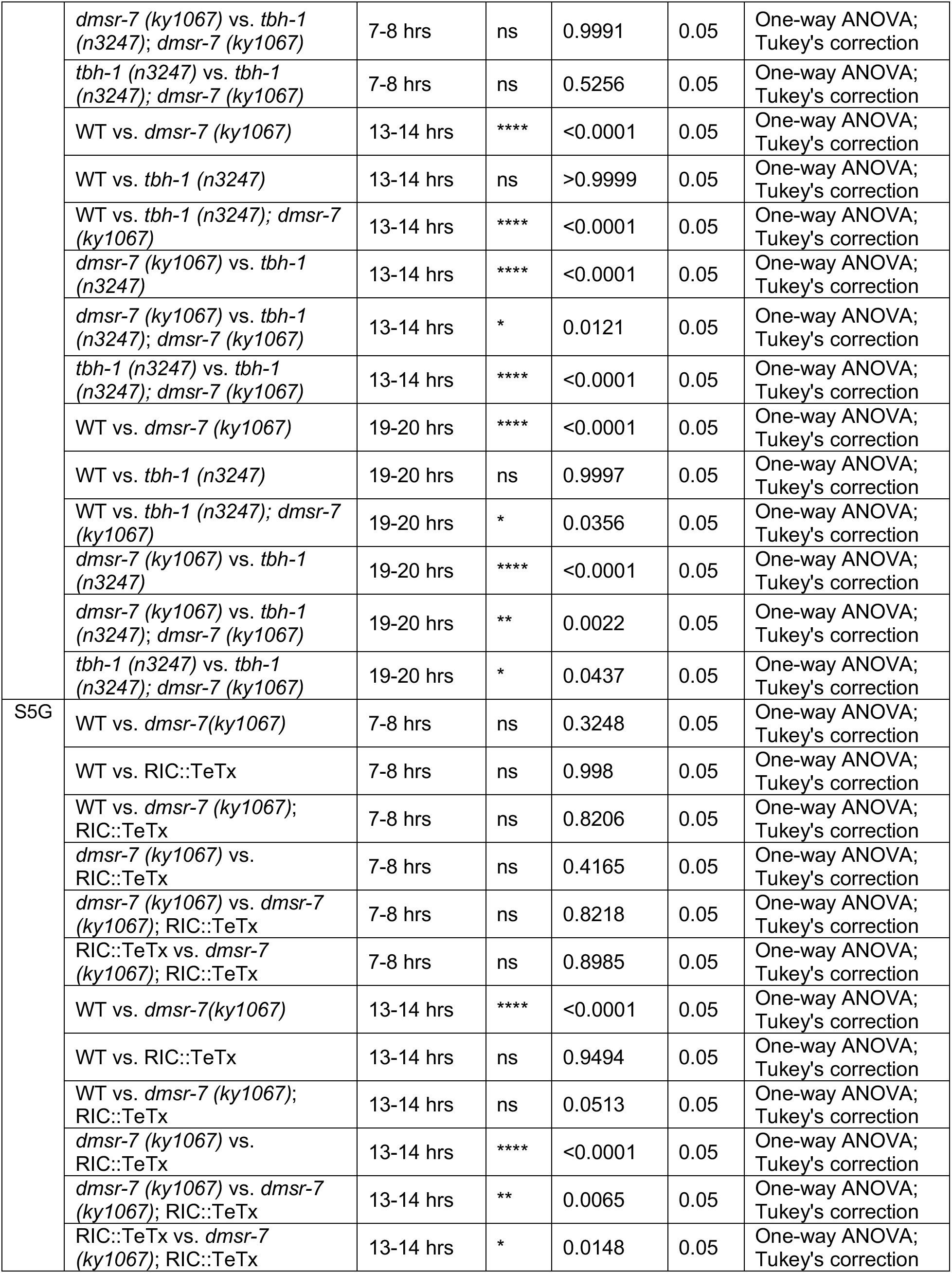
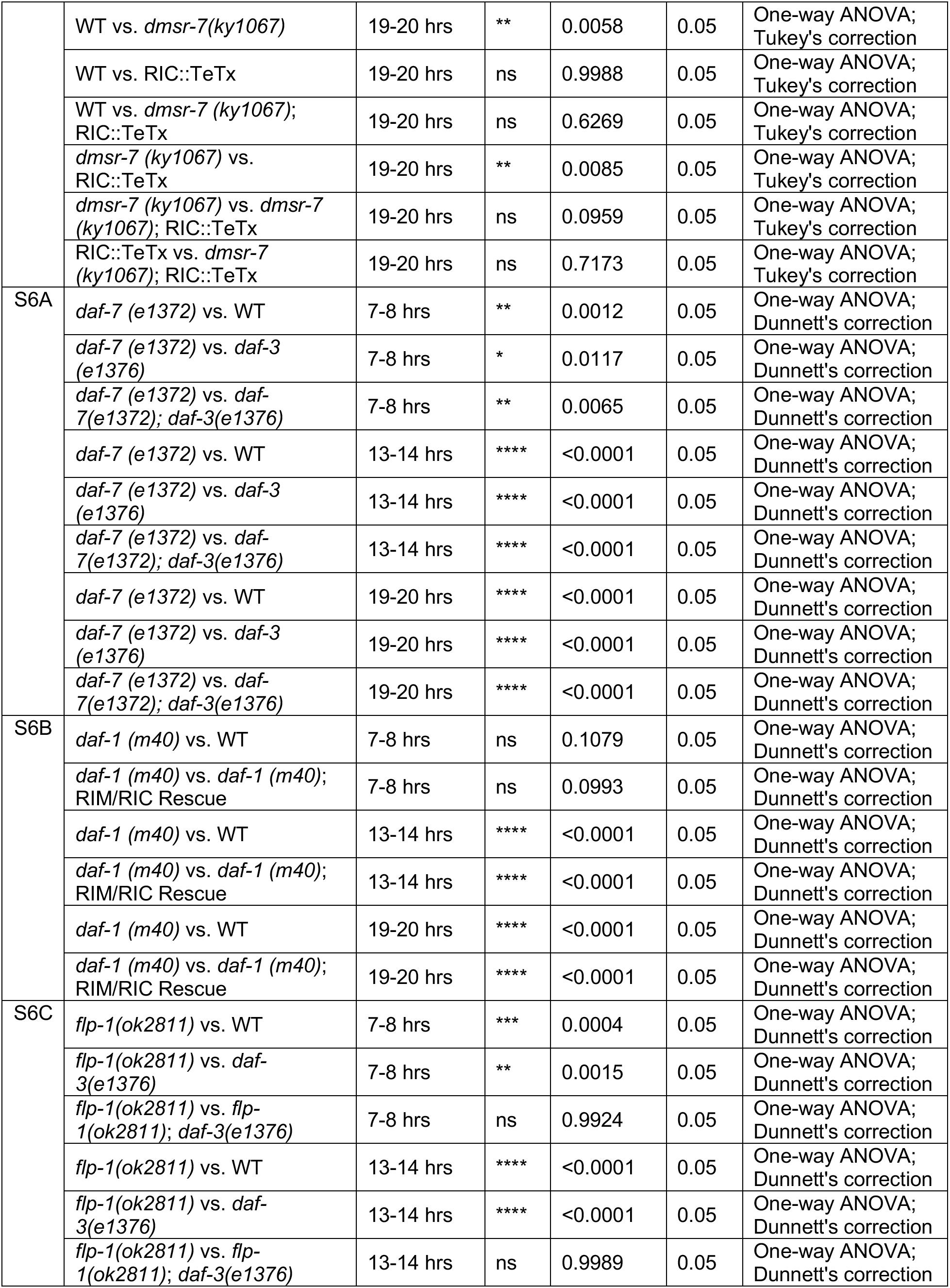
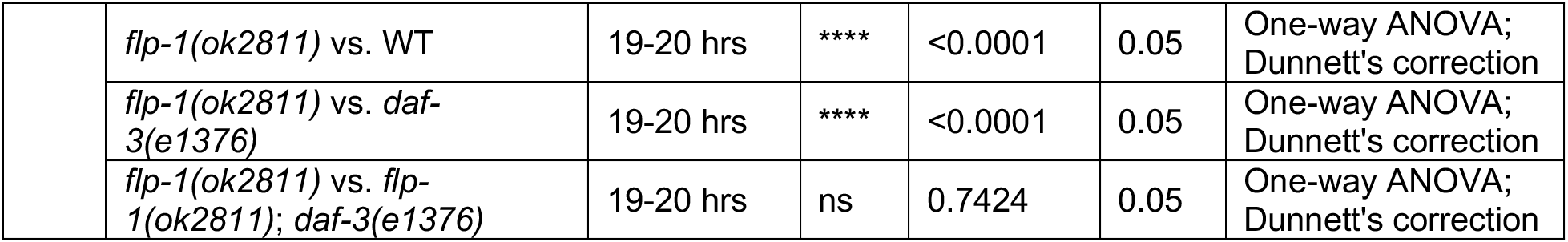
Statistical table.

**Table S5.** Strains generated in this study.

| Name | Strain | Genotype | Relevant Figure(s) |
| --- | --- | --- | --- |
| Wild type | CX0006 | WT |  |
| *egl-3 (n150)* mutant | CX9191 | *egl-3(n150)* V | 1C; S1C-E |
| *egl-3 (ok979)* mutant | VC671 | *egl-3(ok979)* V | 1C |
| *egl-21 (n476)* mutant | KP2018 | *egl-21 (n476)* IV | 1C |
| *egl-21 (n611)* mutant | MT1241 | *egl-21 (n611)* IV | 1C |
| *egl-3* Rescue - Isolate A | CX17653 | *egl-3 (n150)* V, kyEx6219 [*egl-3p::egl-3::SL2::GFP* (5 ng/uL) + *myo-2p::mCherry* (1 ng/uL)] | 1E |
| *egl-3* Rescue - Isolate B | CX17698 | *egl-3 (n150)* V, kyEx6219 [*egl-3p::egl-3::SL2::GFP* (5 ng/uL) + *myo-2p::mCherry* (1 ng/uL)] | 1E |
| *egl-3* Floxed | DCR6978 | *egl-3 (nu1711)* V | 2A, B; S1B-E |
| AVK-specific *egl-3* Knockout [*flp-1p*(513 bp)] - Isolate A | CX18236 | *egl-3 (nu1711)* V; KyEx6532 [*flp-1p*(513 bp)::CRE (20 ng/uL) + *elt-2p::nls::GFP* (2 ng/uL)] | 2B |
| AVK-specific *egl-3* Knockout [*flp-1p*(513 bp)] - Isolate B | CX18237 | *egl-3 (nu1711)* V; KyEx6532 [*flp-1p*(513 bp)::CRE (20 ng/uL) + *elt-2p::nls::GFP* (2 ng/uL)] | 2B |
| AVK::HisCl - Isolate A | CX18087 | kyEx6439 [*flp-1p*(513 bp)::HisCl::SL2::GFP (20ng/uL) + *myo-3p::mCherry* (4ng/uL)] | 2C; S2A |
| AVK::HisCl - Isolate B | CX18088 | kyEx6440 [*flp-1p*(513 bp)::HisCl::SL2::GFP (20ng/uL) + *myo-3p::mCherry* (4ng/uL)] | 2C; S2A |
| AVK::*pkc-1(gf)* - Isolate A | CX18372 | kyEx6595 [*twk-47p::pkc-1(gf)::SL2::mCherry* (30 ng/uL) + *elt-2p::nls-GFP* (2 ng/uL)] | 2D; S2C |
| AVK::*pkc-1(gf)* - Isolate B | CX18373 | kyEx6596 [*twk-47p::pkc-1(gf)::SL2::mCherry* (30 ng/uL) + *elt-2p::nls-GFP* (2 ng/uL)] | 2D; S2C |
| *flp-1* mutant | CX17981 | *flp-1(ok2811)* IV | 2E; 3A, B; S4F; S6C |
| *flp-1* Rescue - Isolate A | CX18062 | *flp-1 (ok2811)* IV; kyEx6423 [*flp-1p*(1.5 kb)::*flp-1::SL2::GFP* (1 ng/uL) + *myo-2p::mCherry* (1 ng/uL)] | 2E; S3C |
| *flp-1* Rescue - Isolate B | CX18063 | *flp-1 (ok2811)* IV; kyEx6423 [*flp-1p*(1.5 kb)::*flp-1::SL2::GFP* (1 ng/uL) + *myo-2p::mCherry* (1 ng/uL)] | 2E |
| *flp-1p::GFP* - Isolate B | CX17931 | kyEx6361 [*flp-1p*(1.5kb)::GFP (20 ng/uL) , *myo-2p::mCherry* (1 ng/uL)] | 3C, E |
| AVK::FLP-1::mCherry Fusion | CX18073 | zxEx825 [*flp-1p*(513 bp)::FLP-1(cDNA)::mCherry + *flp-1p*(513 bp)::GFP] | 3D, F; S3D |
| *dmsr-7* mutant | CX1067 | *dmsr-7* (*ky1067*) V | 4B, D, E; S4E, F; S5B-G |
| *dmsr-7* Rescue - Isolate A | CX18082 | *dmsr-7* (*ky1067*) V; kyEx6434 [*dmsr-7p*::*dmsr-7*::SL2::GFP (10 ng/uL) + *myo-3p*::mCherry (4 ng/uL)] | 4B; S5A |
| *dmsr-7* Rescue - Isolate B | CX18083 | *dmsr-7* (*ky1067*) V; kyEx6434 [*dmsr-7p*::*dmsr-7*::SL2::GFP (10 ng/uL) + *myo-3p*::mCherry (4 ng/uL)] | 4B |
| *dmsr-7* Inverted Rescue | CX18228 | *dmsr-7* (*ky1067*) V; kyEx6524 [*dmsr-7p*::*dmsr-7* Part1::SL2::GFP::*dmsr-7* Part2) (10 ng/uL) + *myo-3p*::mCherry (3 ng/uL)] | 4C, D; S5B-E |
| *dmsr-7* mutant, RIM/RIC Rescue | CX18234 | *dmsr-7* (*ky1067*) V; KyEx6524 [*dmsr-7p*::*dmsr-7* Part1::SL2::GFP::*dmsr-7* Part2) (10 ng/uL) + *myo-3p*::mCherry (3 ng/uL)]; KyEx6531 [*tdc-1p*::CRE (15 ng/uL) + *myo-2p*::mCherry (1 ng/uL)] | 4D |
| *daf-3* (*e1376*) mutant | CB1376 | *daf-3* (*e1376*) X | 4E; S6A, C |
| *dmsr-7* (*ky1067*), *daf-3* (*e1376*) double mutant | CX18455 | *dmsr-7*(*ky1067*) V; *daf-3*(*e1376*) X | 4E |
| RIM/RIC::TeTx | CX14993 | kyEx4962 = *tdc-1p*::TeTx::mCherry (50ng/ul) | 5B |
| *dmsr-7* (*ky1067*); RIM/RIC::TeTx | CX18283 | *dmsr-7*(*ky1067*) V; kyEx4962 [*tdc-1p*::TeTx::mCherry (50ng/ul)] | 5B |
| *tdc-1* (*n3419*) mutant | MT13113 | *tdc-1* (*n3419*) II | 5C |
| *dmsr-7* (*ky1067*); *tdc-1* (*n3419*) double mutant | CX18278 | *tdc-1*(*n3419*) II; *dmsr-7*(*ky1067*) V | 5C |
| RIM/RIC::GFP; *lin-15* (*n765ts*) | QW2 | *zfls1* I [*tdc-1p*::GFP + *lin-15* (*n765ts*) rescue] | 5D, E |
| *dmsr-7* (*ky1067*); RIM/RIC::GFP | CX18456 | *dmsr-7*(*ky1067*) V; *zfls1* I [*tdc-1p*::GFP + *lin-15* (*n765ts*) rescue] | 5D, E |
| Pan-neuronal Knockout of *egl-3* | DCR7147 | *egl-3*(*nu1711*) V; *tmls777* (*rgef-1p*::Cre + *unc-119p*::venus) | S1C |
| Intestinal Knockout of *egl-3* | CX17806 | *egl-3*(*nu1711*) V; *unc-119*(*ed3*) III; *heSi142* [*elt-2p*::nls::CRE + *unc-119* Rescue] X | S1D |
| AVK-specific *egl-3* Knockout (*twk-47p*) | CX18258 | *egl-3*(*nu1711*) V; kyEx6545 [*twk-47p*::CRE (20 ng/uL) + *elt-2p*::nls::GFP (2 ng/uL)] | S1E |
| AVK::TeTx - Isolate A | CX17932 | *kyEx6362 = flp-1p(513 bp)::TeTx (40 ng/uL) , myo-2p::mcherry (1 ng/uL)* | S2B |
| AVK::TeTx - Isolate B | CX17933 | *kyEx6363 = flp-1p(513 bp)::TeTx (40 ng/uL) , myo-2p::mcherry (1 ng/uL)* | S2B |
| *nlp-49 (gk546875)* mutants | CX17982 | *nlp-49 (gk546875) X* | S3A |
| *flp-1 (ok2505)* partial loss of function mutants | CX17980 | *flp-1 (ok2505) IV* | S3B |
| *npr-6 (tm1497); frpr-7 (gk463846)* double mutant | ZX1836 | *npr-6 (tm1497) X; frpr-7 (gk463846) X* | S4A |
| *dmsr-1 (ky1053)* mutant | CX1053 | *dmsr-1 (ky1053) V* | S4B |
| *dmsr-5 (ky1059)* mutant | CX1059 | *dmsr-5 (ky1059) III* | S4C |
| *dmsr-6 (ky1065)* mutant | CX1065 | *dmsr-6 (ky1065) II* | S4D |
| *dmsr-1 (ky1053); dmsr-7(ky1067)* double mutant | CX18072 | *dmsr-1 (ky1053) V; dmsr-7(ky1067) V* | S4E |
| *dmsr-5 (ky1059); dmsr-6 (ky1065); dmsr-7(ky1067)* triple mutant | CX18487 | *dmsr-6 (ky1065) II; dmsr-5 (ky1059) III; dmsr-7(ky1067) V* | S4F |
| *dmsr-7(ky1067)* mutant, Pan-neuronal Rescue | CX18232 | *dmsr-7(ky1067) V; kyEx6524 [dmsr-7p::dmsr-7 Part1::SL2::GFP::dmsr-7 Part2) (10 ng/uL) + myo-3p::mCherry (3 ng/uL)]; KyEx6528 [h20p::CRE (5 ng/uL) + myo-2p::mCherry (1 ng/uL)]* | S5C |
| *dmsr-7(ky1067)* mutant, RIC Rescue | CX18286 | *dmsr-7(ky1067) V; kyEx6524 [dmsr-7p::dmsr-7 Part1::SL2::GFP::dmsr-7 Part2) (10 ng/uL) + myo-3p::mCherry (3 ng/uL)]; KyEx6563 [tbh-1p::CRE (20 ng/uL) + myo-2p::mcherry (1 ng/uL)]* | S5D |
| *dmsr-7(ky1067)* mutant, Cholinergic Neurons Rescue | CX18230 | *dmsr-7(ky1067) V; kyEx6524 [dmsr-7p::dmsr-7 Part1::SL2::GFP::dmsr-7 Part2) (10 ng/uL) + myo-3p::mCherry (3 ng/uL)]; KyEx6526 [unc-17p::CRE (30 ng/uL) + myo-2p::mCherry (1 ng/uL)]* | S5E |
| *tbh-1(n3247)* mutant | MT9455 | *tbh-1 (n3247) X* | S5F |
| *dmsr-7(ky1067); tbh-1(n3247)* double mutant | CX18277 | *dmsr-7(ky1067) V; tbh-1(n3247) X* | S5F |
| RIC::TeTx | CX17912 | *kyEx6345 [tbh-1p::TeTx::mCherry (10 ng/uL) + unc-122p::GFP (5 ng/uL)]* | S5G |
| *dmsr-7* (*ky1067*);<br>RIC::TeTx | CX18488 | *dmsr-7* (*ky1067*) V; *kyEx6345* [*tbh-1p*:TeTx::mCherry (10 ng/uL) + *unc-122p*::GFP (5 ng/uL)] | S5G |
| *daf-7* (*e1372*) mutant | CB1372 | *daf-7* (*e1372*) III | S6A |
| *daf-7* (*e1372*); *daf-3*(*e1376*) double mutant | JT5464 | *daf-7*(*e1372*) III; *daf-3*(*e1376*) X | S6A |
| *daf-1* (*m40*) mutant | DR40 | *daf-1* (*m40*) IV | S6B |
| *daf-1* (*m40*) mutant,<br>RIM/RIC rescue | CX9132 | *daf-1* (*m40*) IV; *kyEx1814* [*tdc-1p*::*daf-1*::SL2::GFP (40 ng/ul) + *elt-2p*::GFP (7 ng/ul)] | S6B |
| *flp-1*(*ok2811*), *daf-3* (*e1376*) double mutant | CX18368 | *flp-1*(*ok2811*) IV; *daf-3* (*e1376*) X | S6C |

